# Epithelial layer unjamming shifts energy metabolism toward glycolysis

**DOI:** 10.1101/2020.06.30.180000

**Authors:** Stephen J. DeCamp, Victor M.K. Tsuda, Jacopo Ferruzzi, Stephan A. Koehler, John T. Giblin, Darren Roblyer, Muhammad H. Zaman, Scott T. Weiss, Margherita DeMarzio, Chan Young Park, Nicolas Chiu Ogassavara, Jennifer Mitchel, James P. Butler, Jeffrey J. Fredberg

## Abstract

In development of an embryo, healing of a wound, or progression of a carcinoma, a requisite event is collective epithelial cellular migration. For example, cells at the advancing front of a wound edge tend to migrate collectively, elongate substantially, and exert tractions more forcefully compared with cells many ranks behind. With regards to energy metabolism, striking spatial gradients have recently been reported in the wounded epithelium, as well as in the tumor, but within the wounded cell layer little is known about the link between mechanical events and underlying energy metabolism. Using the advancing confluent monolayer of MDCKII cells as a model system, here we report at single cell resolution the evolving spatiotemporal fields of cell migration speeds, cell shapes, and traction forces measured simultaneously with fields of multiple indices of cellular energy metabolism. Compared with the epithelial layer that is unwounded, which is non-migratory, solid-like and jammed, the leading edge of the advancing cell layer is shown to become progressively more migratory, fluid-like, and unjammed. In doing so the cytoplasmic redox ratio becomes progressively smaller, the NADH lifetime becomes progressively shorter, and the mitochondrial membrane potential and glucose uptake become progressively larger. These observations indicate that a metabolic shift toward glycolysis accompanies collective cellular migration but show, further, that this shift occurs throughout the cell layer, even in regions where associated changes in cell shapes, traction forces, and migration velocities have yet to penetrate. In characterizing the wound healing process these morphological, mechanical, and metabolic observations, taken on a cell-by-cell basis, comprise the most comprehensive set of biophysical data yet reported. Together, these data suggest the novel hypothesis that the unjammed phase evolved to accommodate fluid-like migratory dynamics during episodes of tissue wound healing, development, and plasticity, but is more energetically expensive compared with the jammed phase, which evolved to maintain a solid-like non-migratory state that is more energetically economical.

**Two sentence summary:** At the leading front of an advancing confluent epithelial layer, each cell tends to migrate, elongate, and pull on its substrate far more than do cells many ranks behind, but little is known about underlying metabolic events. Using the advancing monolayer of MDCKII cells as a model of wound healing, here we show at single cell resolution that physical changes associated with epithelial layer unjamming are accompanied by an overall shift toward glycolytic metabolism.

## INTRODUCTION

From the point of view of energy metabolism, cell migration is costly. Cytoskeletal dynamics generate and transmit the forces that drive cellular migration and, in doing so, can consume an estimated 50% of the cell’s ATP budget^1,2^. As such, a relationship between cellular metabolism and cellular migration is to be expected but the nature of such a relationship has remained unclear. On this topic, much of what is known focuses on cancer biology, the roles of the extracellular matrix (ECM; Supplement 1), and the epithelial-to-mesenchymal transition (EMT). There is consensus that the migrating mesenchymal phenotype exhibits increased aerobic glycolysis^3–8^ and that the Warburg Effect, which is typical of many cancer cells^9,10^, may promote migratory or invasive behavior^11,12^. For example, mesenchymal relative to epithelial prostate cancer cells exhibit increased rates of aerobic glycolysis as well as increased migration speeds and traction forces exerted by each cell on its substrate^4^. Even non-cancerous migratory processes, such as the neural crest cell undergoing EMT, exhibit increased glycolytic flux relative to the non-migratory cell^3^. Similarly, compared to the uninjured epithelial layer, the wounded migrating epithelial cell layer preferentially utilizes glycolysis in much the same manner as migrating mesenchymal cells^13^. Recent spatial imaging of the metabolic state of the wounded epithelium shows large variations in fluorescence lifetimes of NADH^14,15^, an electron carrier molecule ubiquitously involved in energy metabolism. From a mechanistic perspective, cells can modulate their metabolic state in response to extracellular forces through mechanotransduction that is associated with the adhesome and the cytoskeleton^16– 18^, but it remains unclear how metabolic events are connected to collective epithelial cell migration more generally and the cellular unjamming transition (UJT) in particular.

UJT is known to occur during embryonic development^19,20^, injury repair^21^, tumor metastasis^22,23^, and in the asthmatic airways^19,24^. Both the UJT and the EMT potentiate epithelial plasticity and migration, but underlying mechanisms of UJT versus EMT are distinct^19,20^,^25,26^. Unlike the case of the EMT, for example, in the case of the UJT the cell-cell junction and barrier function remain fully intact^25^. The cardinal feature of the UJT, however, is the transition of the confluent epithelial collective from a solid-like, non-migratory, jammed phase to a fluid-like, migratory, unjammed phase^27^. In the jammed epithelium, each cell remains in a fixed spatial relationship to each of its neighbors, as if the collective were frozen. Even though cell-cell junctions remain intact, the UJT enables these cells to unfreeze and thereby more readily swap positions with their neighbors and migrate in fluid-like swirling multicellular packs and eddies. This physical solid-to-fluid transformation is typified by dynamical, morphological, and mechanical alterations^19,24,25,27^. This mechanical plasticity allows for rapid response to tissue perturbations while maintaining uncompromised barrier function^25^. It also suggests the possibility of hijacking or disruption of these processes in epithelial diseases. But little is known about the energetic events that fuel these changes. Data presented here demonstrates that the unjamming cell layer undergoes a dramatic shift in energy metabolism toward glycolysis.

## RESULTS

We utilized multiple modes of metabolic imaging simultaneously with measurements of cell mechanics, dynamics, and morphology. These measurements, taken together, reveal the complex spatiotemporal mechano-bioenergetic changes that occur in the confluent epithelial layer both while confined by a barrier and while undergoing collective migration into free space. We used a removable polydimethyl-siloxane (PDMS) mask on polyacrylamide (PA) gel to grow an epithelial cell monolayer under geometric confinement (Fig. 1a,b). The PA gel was 100 μm thick, had a shear modulus of 9.6 kPa, and was coated with collagen-I (Methods). Just below its apical surface this gel also contained many fiducial markers comprised of fluorescent labeled beads (0.2 µm diameter) whose displacements in response to cellular contractile forces were used for traction microscopy^28^. This gel was then seeded with Madin-Darby Canine Kidney (MDCKII) cells which were grown to confluence until they tiled the entire available surface (Fig. 1c). After the cell layer reached confluence, the PDMS barrier was removed and, in response, the cell layer migrated into the newly created free space (Fig. 1d,e). As has been characterized elsewhere, this simplified model recapitulates cardinal features of epithelial wound healing but allows for measurement of cell mechanics, dynamics, and morphology^29^. In order to measure cell metabolic properties simultaneously with mechanics, these MDCKII cells were stably transfected with mCherry-Peredox-NLS (Methods) which enabled assessment of the cytoplasmic redox potential (NAD+/NADH) across the cell layer with single cell resolution^30^ (Fig. 1f–j, Supplementary Fig. 1). Additionally, we used fluorescence lifetime imaging (FLIM) to isolate contributions from free and enzyme bound NADH across the cell layer. To further map metabolic heterogeneities developing during collective migration, we quantified the total glucose uptake per cell using the 2-NBDG glucose uptake assay and measured the mitochondrial membrane potential using a membrane potential dependent dye, TMRE (Methods).

**Figure 1:**
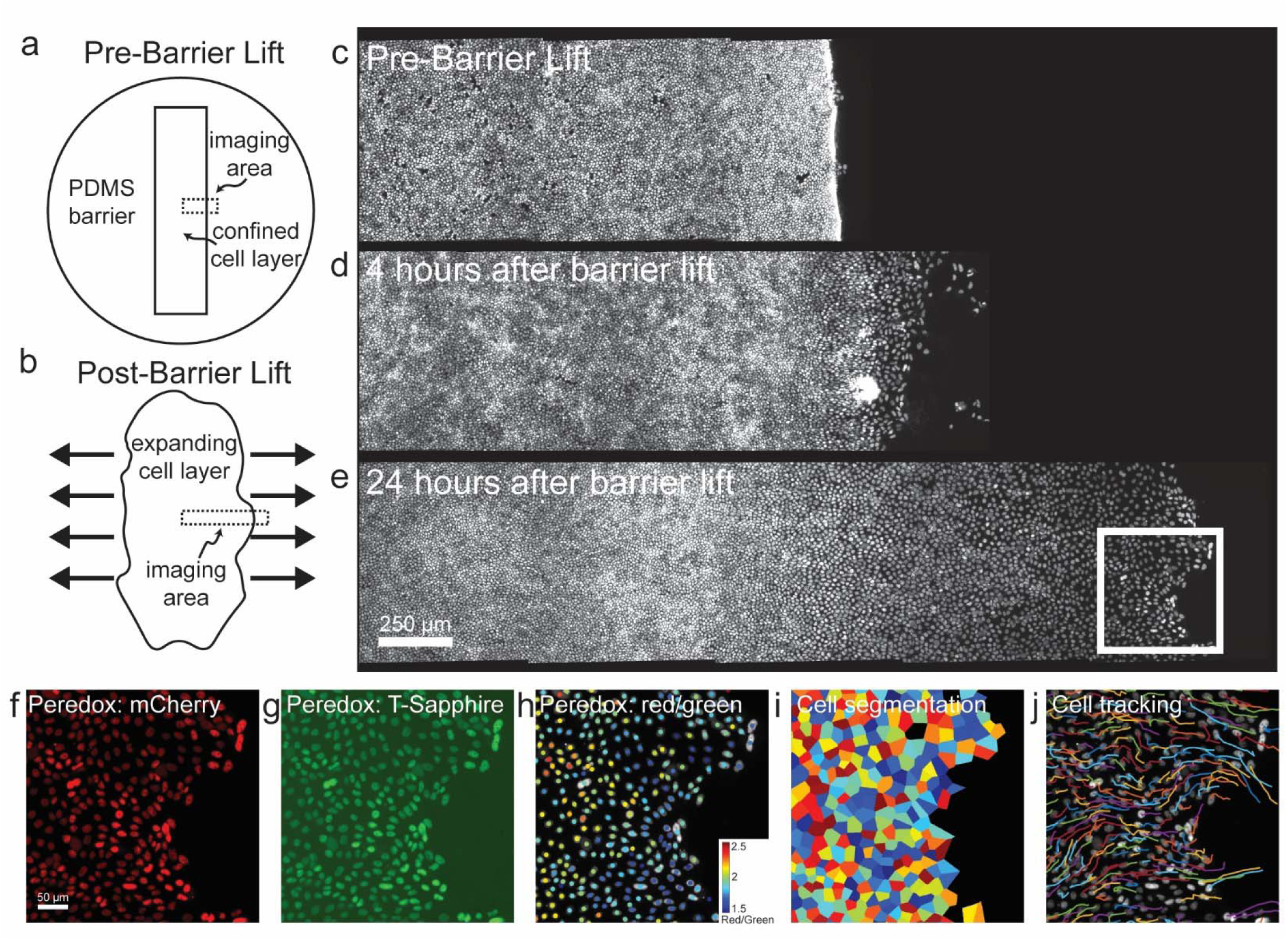
Sudden creation of free space launches gradients of cell morphology, migration, and energy metabolism. **a** Schematic of the MDCKII cell layer initially confined by a rectangular PDMS barrier. **b** When the PDMS barrier is removed, free space is created and cells in the layer migrate into free space and the layer expands as indicated by the bold black arrows. We image the cells in a region across the cell layer as indicated by the dashed rectangular outline. **c-e** Fluorescent microscopy images of the mCherry-Peredox labeled cell nuclei before PDMS barrier lift and at 4 hrs and 24 hrs after barrier removal. The center position of the expanding cell layers are aligned at the left hand side of the X-axis and with x-positions increasing towards the right. A black background has been added to frame the images. This MDCKII cell line expresses the Peredox-NLS fluorescence biosensor that reports the cytoplasmic redox potential. Here, we show a zoomed-in region of interest outlined by the white box in panel **e**, which depicts the leading edge of the advancing cell layer. **f** The mCherry (red) channel of the Peredox sensor is used for normalization of variations in sensor expression. **g** The T-Sapphire (green) channel is NADH-sensitive. **h** The ratio of the red-to-green channel is proportional to the cytoplasmic NAD+/NADH ratio. Here, we measure the fluorescence ratio with single-cell resolution. The color coded dots are centered over each identified nucleus in the image and denote the ratio of the two fluorescence channels. **i** In addition to cell redox, cell morphology is measured based by tessellating nuclear centroids. **j** Cell dynamics are measured by tracking nuclei over short time-lapse movies. Shown here are tracks of cells migrating from left-to-right into free space. Each track represents the successive position over two-hours of imaging.

### Redox ratios decrease as cells unjam and become migratory

When confined by the PDMS mask, cellular speeds remained small and uniform (Fig. 2a top panel). With the exception of an edge artifact near the PDMS mask, traction forces exerted throughout the layer, cell area, and cell shape (as reflected in the aspect ratio) all remained small and uniform (Fig. 2b,c,d top panels; Supplementary Movie 1). We found that the redox ratio measured by the Peredox sensor remained high and showed little variation across the cell layer, thus suggesting that the cell layer while initially confined by the PDMS barrier is initially in a homogenous metabolic state (Fig. 2e,f). When the PDMS mask was lifted, however, free space was created into which the monolayer could migrate (Supplementary Movie 2). At the earlier time point (4 hr), the monolayer front had advanced by approximately 500 μm. As reported previously, cells at the advancing front increased in aspect ratio as they advanced, and elongated in the direction of migration (Fig. 2c middle panel, Supplementary Movie 2)^29^. In this region cell speeds and traction forces increased while, in contrast, cells deep in the center of the layer showed little change in these variables (Fig. 2a,b middle panels). Relative to cells in the layer center, the redox ratio at the advancing layer edge at the 4 hr timepoint decreased substantially (Fig. 2e,f). At the later time point (24 hr), the front had advanced by approximately 1.3 mm. By this later time point, perturbations in cell shapes, cell area, migrations speeds, traction forces, and cell redox ratio had propagated within the cell layer substantial distances retrograde (Fig. 2a–e bottom panels). By this later timepoint, the cytoplasmic redox ratio established across the cell layer a steep gradient that spanned nearly an order of magnitude (Fig. 2e,f). These measurements remain within the range reported in the literature for cells maintained in normal or nutrient starved conditions; typical cytoplasmic NAD+/NADH ratios range from 10 to 1000^30,31^. Furthermore, the low cytoplasmic NAD+/NADH ratio near the migrating edge is consistent with a shift towards glycolytic activity opposing LDH^30^. Although Serra-Picamal *et al*. measured no metabolic indices, they reported that creation of a free space launches not only an advancing wave of migration, which acts to fill that free space and thus heal the wound, but also a retrograde wave of unjamming that acts to mobilize cells in the ranks behind and thereby recruit them to the advancing front^29,32,33^. In this unjamming process retrograde waves of cell deformation trigger retrograde waves of ERK activation in a sustained mechano-chemical feedback loop^34^. These changes in mechanical, chemical, and morphological indices spatially coincide with the regions in the epithelial cell layer where the redox ratio is dramatically reduced (Fig. 2a–e).

**Figure 2:**
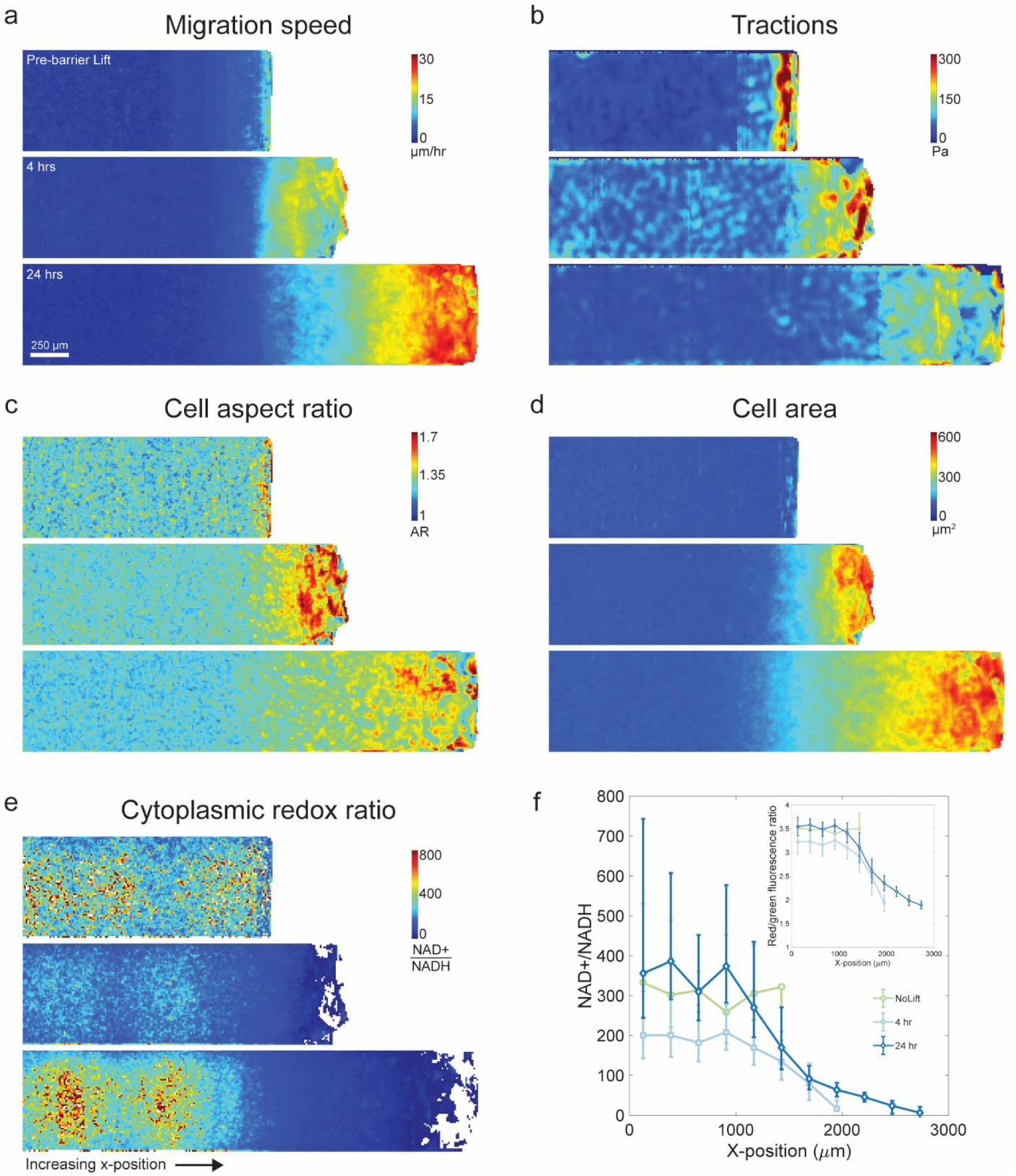
Cell redox potential decreases upon epithelial layer unjamming. **Top Panels:** Prior to lifting the PDMS barrier, cells are confined and jammed. **a** The cell migration speed shows that very little migration is occurring in the layer. **b** Traction forces are low throughout the layer but show an increased edge-effect near the PDMS barrier. **c,d** Cells are uniformly small in area and round in shape. **e,f** The cytoplasmic redox potential (NAD+/NADH) measured via the Peredox biosensor is high throughout the cell layer. **Middle Panels:** 4 hrs after lifting the PDMS barrier, cells near the layer edge begin to migrate into the free space and the layer expands. Cell migration speeds are increased near the advancing edge. Traction forces are elevated throughout the layer and a steep gradient appears from the leading edge into the bulk. Cell area is dramatically increased and cell shapes become elongated near the advancing edge. The cell redox potential decreases at the leading edge. **Bottom Panels:** 24 hrs after lifting the PDMS barrier, the layer has expanded to nearly twice the extent of the confined layer. Migration speeds at the leading edge continue to increase as the cell layer migrates into the free space. Traction forces are elevated at the migrating front and cells have substantially expanded in area and elongated in shape. The cytoplasmic redox potential in the migrating cells remains low relative to the jammed cells near the center of the layer. Error bars represent the standard deviation of the mean of the fluorescence ratio which are subsequently transformed to a redox potential using the Peredox fluorescence response curve (Supplementary Fig 1). As the calibration is non-linear, the conversion results in asymmetric error bars.

Tendencies noted above, as well systematic relationships between morphological, migratory, mechanical and metabolic indices, are analyzed statistically and quantified on a cell-by-cell basis in Supplement 3. These analyses indicate that among all variables measured, the strongest statistical predictor of local migration speed was local cell perimeter, but with an important contribution from the NAD+/NADH ratio. Similarly, the strongest statistical predictor of the local NAD+/NADH ratio was also local cell perimeter.

### NADH lifetimes decrease at the migrating edge and in the non-migratory bulk

To confirm the spatial shifts in redox state of the cellular collective, we tracked NADH lifetimes across the expanding monolayer using FLIM. NADH is autofluorescent and the time it remains in an excited state before emitting a photon, known as the NADH lifetime, is sensitive to its binding state to metabolic enzymes^35,36^. Multiphoton excitation of NADH, measurement of the photon times of arrival^37,38^, and double exponential fitting of the resulting temporal distributions allow us to quantify the mean lifetime and separate the fraction of free and enzyme-bound NADH (Methods). Shifts in NADH lifetimes and in the ratio of free-to-bound NADH are known to reflect the redox state of epithelial cells^39^ and respond to inhibition of different metabolic pathways both *in vitro and in vivo*^40–42^.

Prior to barrier lifting and after barrier lifting at 4 hrs and 24 hrs, non-migrating cells in the confined jammed layer exhibited a uniform and elevated NADH mean lifetime (Fig. 3a,b) indicative of a larger fraction of NADH in the bound-state (Fig. 3c,d). Relative to before barrier lift, at 4 hrs the mean NADH lifetime was slightly reduced across the layer (Fig. 3a,b) and the free-to-bound ratio increased (Fig. 3c,d). At 24 hrs after lifting the barrier, a spatial gradient emerged whereby cells at the migrating unjammed front showed a sharp decrease in the mean NADH lifetime (Fig. 3a,b) and an increase in the free-to-bound ratio (Fig. 3c,d) with respect to the jammed bulk. Surprisingly, soon after barrier lifting, cells in the non-migratory and jammed center of the layer also exhibited a marked decrease in mean NADH lifetimes. Together with the results from the Peredox bioprobe, these FLIM results are consistent with an overall shift in the unjamming epithelial layer towards glycolysis^40,42–44^.

**Figure 3:**
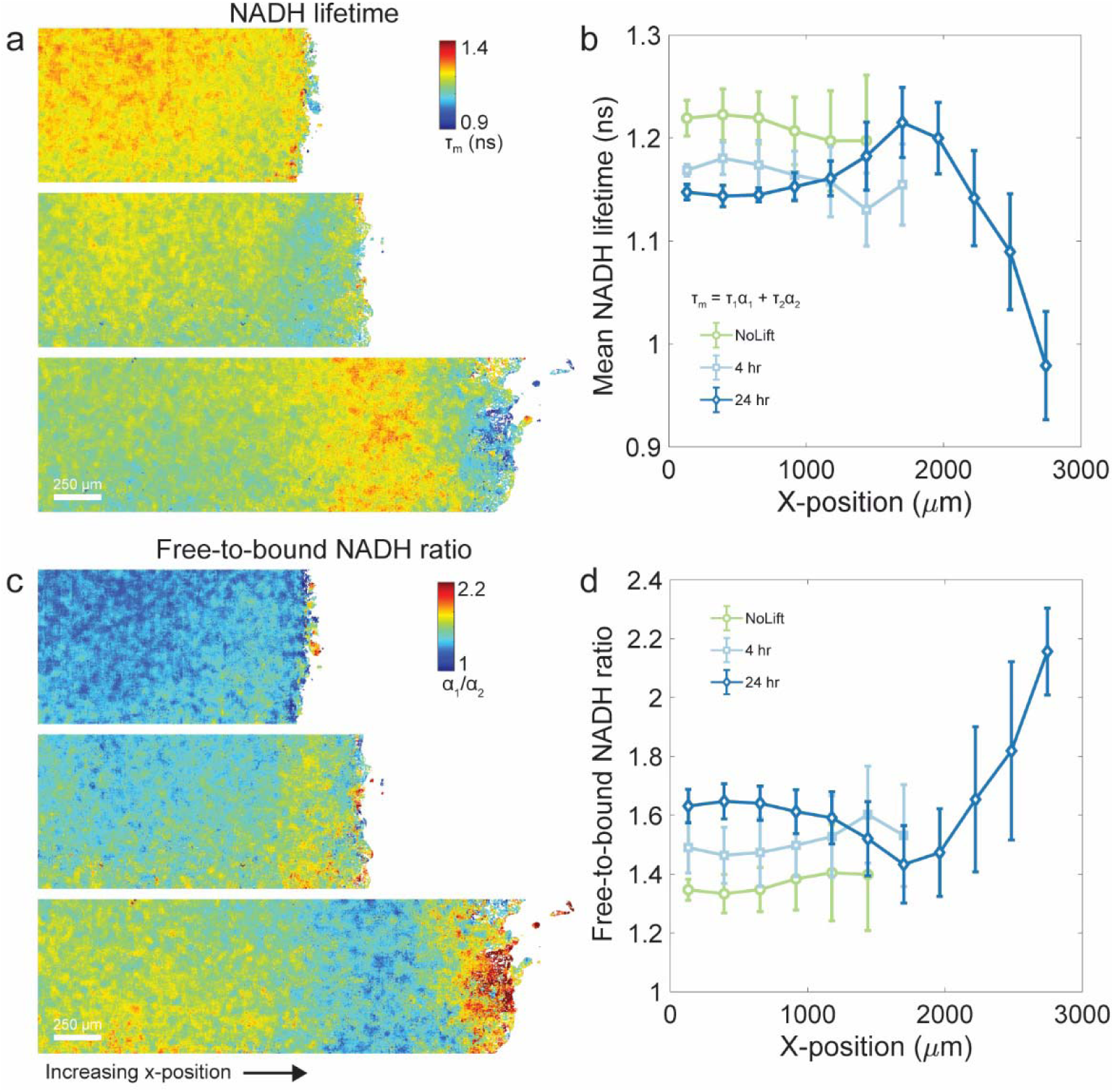
Fluorescence Lifetime Imaging Microscopy (FLIM) reveals that a shift toward glycolytic metabolism occurs in the unjamming cell layer. **a** Composite color maps of the mean NADH lifetime in the non-migrating cell layer confined by the PDMS barrier and at timepoints 4 hrs and 24 hrs after barrier removal. Prior to barrier lifting, cells have a uniform mean NADH lifetime throughout the layer. At 4 hrs after barrier lifting, the layer begins to expand and the mean NADH lifetime is reduced. After 24 hrs, a complex spatial gradient emerges across the expanding cell layer. Cells at the unjammed and advancing front of the cell layer have a drastically reduced mean NADH lifetime relative to cells in the non-migrating layer. Similarly, cells in the jammed bulk show a decreased mean NADH lifetime relative to the pre-barrier and 4 hr timepoints. **b** We plot traces of the mean NADH lifetime prior to barrier lift and at 4 hrs and 24 hrs after barrier lift. The data is averaged along the entire y-axis and into bins along the x-axis. Error bars show layer-to-layer standard deviations of the mean NADH lifetimes. **c** Composite colormaps of the ratio of free to bound NADH in the non-migrating cell layer confined by the PDMS barrier and at timepoints 4 hrs and 24 hrs after barrier removal. Prior to barrier lift, the free-to-bound NADH ratio is uniform and low throughout the cell layer. At 4 hrs after barrier lifting, the layer shifts towards a higher fraction of free NADH. After 24 hrs, cells near the leading edge as well as cells in the bulk have shifted significantly towards a high fraction of free NADH. **d** We plot traces of the fraction of free-to-bound NADH prior to barrier lift and at 4 hrs and 24 hrs after barrier lift. The data is averaged along the entire y-axis and into bins along the x-axis. Error bars show layer-to-layer standard deviations of the free-to-bound NADH ratio.

### Glucose uptake increases in the migrating edge and in the non-migratory bulk

To examine more specifically cell glycolytic activity, we then mapped glucose uptake in wild-type MDCKII cells using 2-NBDG, a fluorescent, non-catabolizable glucose analog (Methods)^45–47^. In the non-migratory and confined layer prior to barrier lifting, cells throughout the layer showed uniform and small glucose uptake per cell, i.e., after normalization by cell area (Fig. 4a,b). At timepoints after lifting the barrier, glucose uptake per cell area progressively increased everywhere throughout the layer, with the largest increase occurring in cells near the leading edge (Fig. 4a,b). This glucose uptake profile across the unjamming cell layer is consistent with the trends in the NADH lifetimes measured by FLIM. Further, it confirms that a metabolic shift toward glycolysis occurred in regions deep within the layer, beyond those regions where cells had unjammed, mobilized, and generated increased tractions.

**Figure 4:**
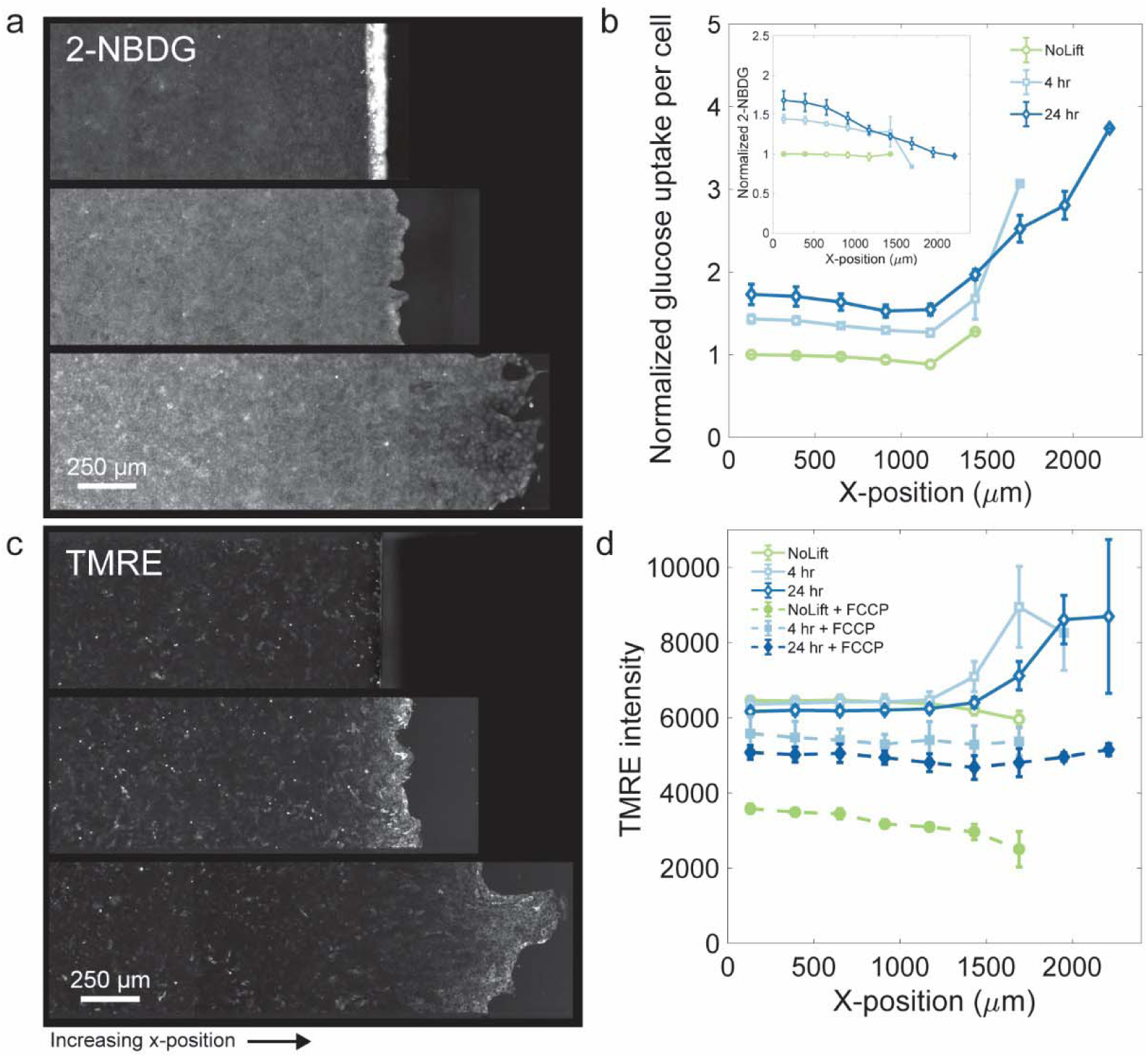
Glucose uptake per cell increases everywhere while mitochondrial membrane potential increases only near the advancing edge. To investigate the extent to which cells within the layer shift toward glycolysis, we employ the glucose uptake assay by imaging the epithelial cell layer after incubation with 2-NBDG, a fluorescent non-catabolizable glucose analog that accumulates within cells after uptake. **a** Representative fluorescence images of the 2-NBDG signal are shown at timepoints prior to barrier lifting and at 4 hrs and 24 hrs after barrier lifting. Cells confined by the PDMS barrier have a uniform and low 2-NBDG intensity throughout the layer. The PDMS barrier appears bright in the image as 2-NBDG tends to non-specifically adhere to the cut PDMS edge. At 4 hrs and 24 hrs after barrier lifting, the 2-NBDG intensity progressively increases in the jammed interior of the cell layer and remains low at the advancing edge. **b** We normalize the 2-NBDG fluorescence intensity of all three timepoints relative to the 2-NBDG intensity at the center of the confined cell layer of the pre-barrier lift condition (inset). Additionally, the average 2-NDBG fluorescence intensity across the cell layer is normalized by the average cell area measurements from tessellations of the Peredox cell nucleus positions. This reveals that on a per-cell basis, prior to barrier lifting, the 2-NBDG intensity is uniform and low. At timepoints after barrier lifting, 2-NDBG uptake per cell, relative to the confined layer, becomes progressively elevated. The 2-NBDG intensity is dramatically increased near the advancing front. To investigate the mitochondrial membrane potential in cells throughout the epithelial cell layer, we use TMRE, a membrane potential dependent dye, imaged on a fluorescence microscope. **c** Representative fluorescence images of the TMRE intensity are shown at timepoints prior to barrier lifting and at 4 hrs and 24 hrs after barrier lifting. Prior to barrier lifting, TMRE fluorescence intensity across the layer remains low and uniform. At 4 hrs after barrier lifting, the TMRE signal remains low in the jammed bulk but becomes more intense near the migrating edge. After 24 hrs, the TMRE signal remains elevated only in the cells at and immediately behind the leading edge. **d** Composite traces of the TMRE image intensity for data prior to barrier lift and at 4 hrs and 24 hrs after barrier lift. TMRE intensity is observed to increase after barrier lift near the leading edge indicating an increased mitochondrial membrane potential in migrating cells, but remains low everywhere else. Cell layers treated with FCCP show a significant reduction in TMRE signal, indicating a drop in mitochondrial membrane potential as expected. Interestingly, the drop in the TMRE signal in the confined and jammed layer is significantly larger than the drop in the migrating cell layers.

### Mitochondria membrane potential increases with cell migration, but only near the leading edge

Mitochondrial membrane potential (MMP) drives ATP production by powering ATP synthase. As such, MMP is a key bioenergetic quantity related to ATP flux during cell respiration. To capture spatiotemporal changes in MMP across the cell layer, we used the lipophilic cationic dye, Tetramethylrhodamine ethyl ester perchlorate (TMRE) (Methods)^48^. TMRE fluorescence intensity across the confined, immotile, and jammed epithelial layer was small and uniform (Fig. 4c,d). At 4 hrs and 24 hrs after the barrier was lifted, only the cells within a few cell-lengths of the migrating edge displayed increased TMRE fluorescence signal (Fig. 4c,d). When we short-circuited the mitochondrial membrane by treating the cells with the mitochondrial membrane potential decoupler, FCCP, TMRE fluorescence signals were substantially reduced (Supplementary Fig. 4). To test if the variation of TMRE fluorescence was related to mitochondrial mass, we stained mitochondria with MitoView-green, a membrane potential independent dye. With the exception of first row leader cells, the MitoView signal was essentially uniform throughout the layer (Supplementary Fig. 4), thereby suggesting that MMP, as indicated by TMRE, is moderately increased at the advancing edge but remains low everywhere else in the layer.

## DISCUSSION

The epithelial layer is ordinarily non-migratory, jammed and quiescent but all the time retains the capacity to unjam and, in doing so, transition to a migratory phase in response to the demand for tissue development, growth, remodeling or wound repair^19,29,49^. It has been suggested that these salutary unjamming behaviors can sometimes become hijacked in disease, however, and thereby initiate unwelcomed cellular migration and tissue remodeling as in the cases of tumor invasion or asthmatic remodeling of the airway wall^24,50–53^. In the advancing confluent MDCKII monolayer, here we have integrated multiple modes of metabolic imaging in order to characterize with single cell precision the metabolic state simultaneously with metrics of cell morphology, migration, and mechanics. The jammed non-migratory epithelial layer confined by the PDMS barrier exhibits small and uniform cell speeds, traction forces, cell areas, and aspect ratios. The confined cell layer also exhibits a homogenous metabolic state as evidenced by the uniform cytoplasmic NAD+/NADH ratio, NADH lifetime, glucose uptake, and MMP.

After lifting the barrier, cells at the leading edge unjam whereby they exert higher traction forces, elongate in shape, and migrate faster. At the unjammed, migrating edge cells exhibit decreased cytoplasmic NAD+/NADH ratio, reduced NADH lifetime, and increased glucose uptake (Fig. 5). These results, taken together, suggest that the UJT provokes a shift toward glycolytic energy metabolism. What drives these complex spatiotemporal gradients at the migrating front? Cells migrating into free space at the leading edge of an advancing cell layer are known to undergo cell cycle reentry^54^. Proliferative cells thus require additional glycolytic activity to fuel growth processes^^10,55^^. On the other hand, cytoskeletal restructuring, which is necessary to support force transduction through the epithelial cell layer and to generate traction for collective cell migration, is associated with a shift toward glycolysis^8,18^ in order to fuel this energetic demand^1,2,56^. An unanswered question therefore becomes the extent to which metabolic shifts are driven by the variable cellular activities that occur near the leading edge of a migrating cell layer, and how the metabolic budget is divided amongst these tasks.

**Figure 5:**
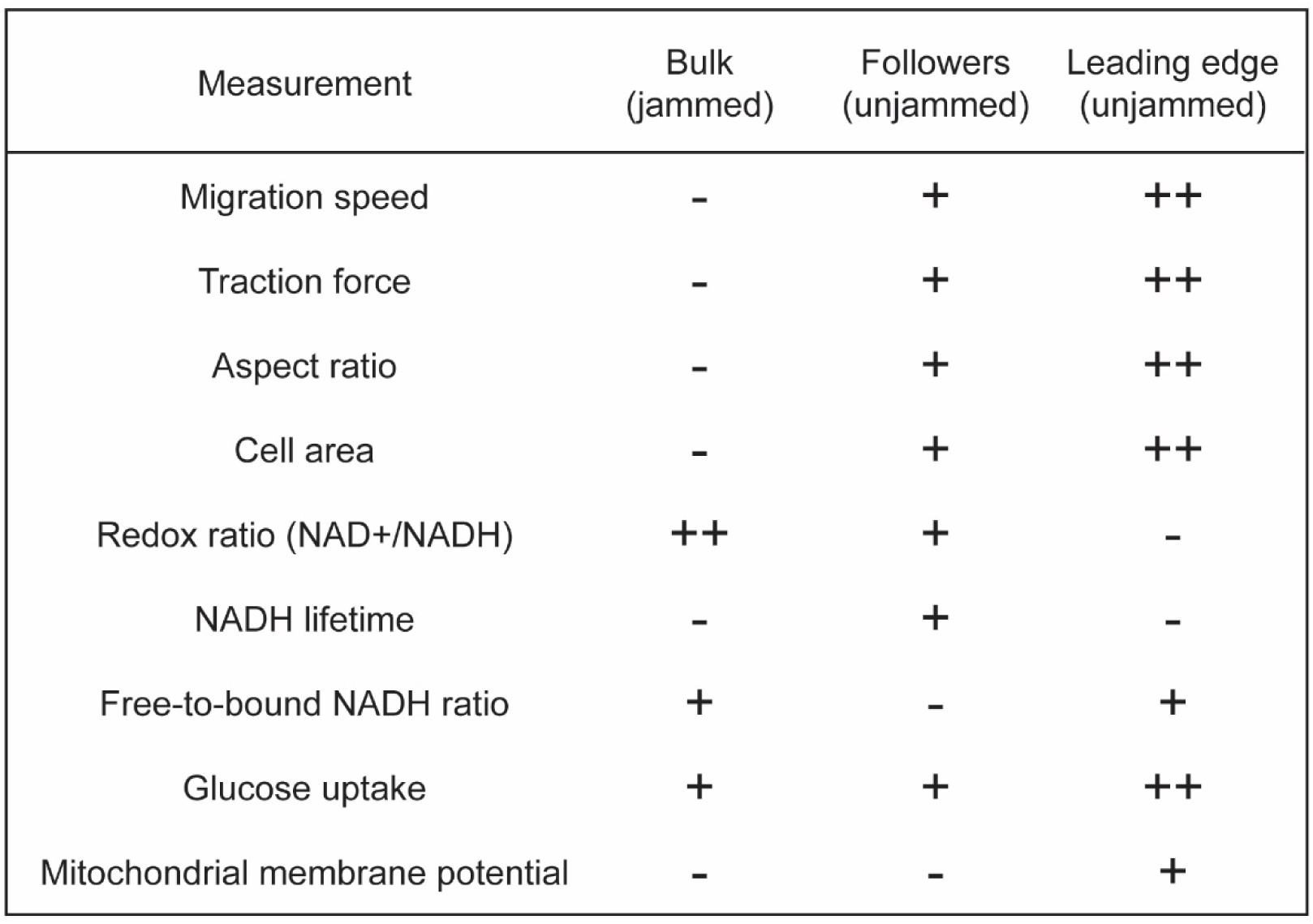
Summary of results. To summarize our findings, we define three regions of interest in which distinctive mechanical and metabolic changes occur in the cell layer; areas which remain non-migratory and jammed in bulk far from the leading edge, unjammed and migratory cells immediately behind the leader cells, and cells at the unjammed and migratory leading edge of the cell layer. For each of the mechanical, dynamical, morphological, and metabolic indices that we have measured, we indicate the qualitative change that occurs in each region after lifting the barrier.

Far from the leading edge cells tend to exhibit progressively slower migration, smaller aspect ratios, smaller tractions, and closer proximity to a jamming transition (Fig. 5)^19,24,27^. Surprisingly, metabolic gradients arose after barrier lifting - even into these regions far from the leading edge where changes of cell shapes, traction forces, and migration speeds had yet to appear. In these regions, jammed cells far from the advancing edge underwent a marked decrease in NADH lifetime (Fig. 3) and an increase in glucose uptake (Fig. 4), consistent with a shift toward glycolysis. In contrast to the cells at the leading edge, however, in the jammed bulk, the cytoplasmic NAD+/NADH remained elevated. Our finding of an elevated MMP at the leading edge and a low MMP in the jammed bulk suggests that the regional variation in the NAD+/NADH signal may be accounted for by LDH activity and mitochondrial demand for pyruvate. Inhibition of LDH and the diversion of carbon to the TCA cycle have been shown to lead to dramatic shifts in NADH lifetimes as measured by FLIM^42,57^. Such a mechanism can maintain the oxidized NADH-NAD+ state in the jammed bulk and is still consistent with an overall shift toward glycolysis.

What drives the metabolic gradient in the jammed non-migratory bulk? In cells grown on PA gels, long distance substrate-deformation fields caused by cell-cell interactions during collective migration can drive large-scale and long-distance correlations in cell motion, particularly in regions of high cell density as is found far from the advancing edge^58^. These deformation fields have been shown to induce mechanical cooperativity throughout the cell layer. As such, we hypothesize that the long-distance transmission of extracellular forces through the elastic substrate, generated by collective cell migration near the advancing front, has the potential to induce glycolysis, even far back from the migrating edge, through the stimulation of cytoskeletal-based mechanotransductive pathways such as PI3-K mediated RAC activation found in MCF10A cells^18^, or LKB1 mediated AMPK activation as found in MCF10A and MDCK cells^16^. While mechanical force transmission through the cell layer is likely to affect the metabolic state of those cells, the degree to which biochemical signaling and mechanical perturbations respectively induce a metabolic shift in the unjamming cell layer remains an open question.

Taken together, these results define the spatiotemporal metabolic signature that accompanies UJT. These new observations link on a regional basis many of the multifaceted events –morphological, mechanical, and metabolic– that play out in the wound healing process. However, these observations are subject to several caveats and limitations. For instance, common techniques such as the Seahorse FX Analyzer measure metabolic flux (extracellular acidification rate or oxygen consumption rate) but do not provide spatial resolution necessary to map local cell mechanics with concurrent metabolic changes. As such, we chose to measure a series of metabolic indices that could be imaged with spatial resolution which, when integrated, reveal changes to major metabolic pathways. The increase in MMP near the leading edge along with the shift towards glycolysis and overall increase in glucose uptake throughout the layer is highly suggestive of an increase in metabolic demand, but the metabolic flux through any specific pathway remains unclear. It stands as a challenge to the field to further integrate metabolic rate information with mechanical and dynamical data in order to better understand metabolic budgets associated with cell migration on a per-cell basis within confluent tissues. With regards to the methods we employ here, the Peredox bioprobe is limited in efficacy to the cytoplasm, which serves as both a strength and a weakness in selectively assessing sub-compartmentalized metabolic processes. Additionally, in some cases where key metabolic reactions are subject to far-from-equilibrium conditions, measurements of NAD+/NADH that utilize lactate/pyruvate titration^31^ may incorrectly estimate the redox ratio^59^. Regarding FLIM, the autofluorescence spectrum of NADH and NADPH are indistinguishable. Even though the NAD pool is typically much larger than the NADP pool, the fraction of the reduced forms are often of similar magnitude^60^. As such, while we focus primarily on the NAD pool in our discussion as it pertains to energy metabolism, we cannot rule out the role of NADPH in anabolic processes as contributing to these metabolic gradients. Lastly, while the overall change in the NADH lifetime signal is consistent with a shift towards glycolytic activity, we find that a biphasic trend emerges at the 24 hr timepoint, which suggests that additional metabolic processes beyond what we explore here may be at play in the advancing cell sheet. These limitations stress the importance of integrating multiple modalities of information relevant to cell metabolic processes to avoid pitfalls in measurement error and to maintain check-and-balances in signal interpretation.

As regards molecular mechanisms of unjamming, our laboratory has recently reported results using a different epithelial system^26,61^. In the case of human bronchial epithelial cells, genome-wide RNA-Seq, protein-protein Interaction analysis, and gene ontology analysis of temporal networks show that epithelial cell unjamming involves the interplay of downstream signaling pathways involving integrins, ERK and JNK. In that system, moreover, gene ontology analyses on differentially expressed genes at early time points identify enrichment of the lipid metabolic process, the cellular ketone metabolic process, regulation of small molecule metabolic processes, and regulation of cellular carbohydrate metabolic process. These processes involve oxidization of fatty acids and are subsequently metabolized in mitochondria for ATP/ energy generation. Additionally, the expression of the mitochondrial carrier protein SLC25A25 involved in uptake and efflux of adenine nucleotides into or from the mitochondria, was significantly induced and hint toward a high energy need and production in the initial phase of UJT that is covered by fatty acid oxidization. In unjammed and migrating cells, the use of unsaturated fatty acid metabolic process to generate energy and cell communication by electrical coupling were significantly enriched. In addition, these unjammed epithelial cells induced the expression of the glucose transporter (SLC2A6) to allow higher glucose uptake from the environment for energy production. To meet the energy need of unjammed migrating cells in that system, these observations, taken together, suggest that the unjamming response initially employs fatty acid oxidation and later enhanced glucose uptake.

Based upon cell-by-cell statistical analysis as shown in Supplement 3, an empirical relationship emerged linking local redox ratio mainly to cell perimeter. This relationship calls to mind how integrins and cadherins interact in a force-dependent manner with a master regulator of energy homeostasis, namely, AMP-activated protein kinase (AMPK) and its upstream activator, Liver Kinase B1 (LKB1)^62^. Accordingly, a plausible mechanistic explanation of this empirical relationship between cell morphology and cell energy metabolism is that local integrin-dependent traction forces generate greater cadherin-dependent intercellular forces^28,63^. These forces lead to more cell stretch and thus additional cell perimeter. Through this mechanism, it is therefore possible that the large cell perimeter leads to a greater extent of glycolytic activation. As the cytoplasmic redox ratio is sensitive to both glycolytic and mitochondrial activity, there exists an alternative notion linking the redox ratio to cell perimeter. Repeated stretching has been shown to regulate ATP synthase and mitochondrial ATP production^17^. As such, it is possible that cells with excessive perimeter, indicative of cell-stretch, could exhibit an altered cellular redox ratio through this mechanism.

### Epithelial unjamming is energetically expensive whereas jamming is economical

The epithelial layer is understood to be a highly dynamic tissue. Here we show this statement to be true but incomplete. As a result of the interconversion between jammed versus unjammed phases, cellular dynamics in the very same tissue can differ dramatically in space and time. The jammed phase of the confluent epithelial collective is solid-like and non-migratory^19,21,24,28,64^. The unjammed phase of the confluent epithelial collective is fluid-like, migratory, and shifted toward glycolytic energy metabolism. This metabolic shift suggests that the unjammed layer possesses an increased metabolic demand that is fulfilled by the fast but inefficient glycolytic energy metabolism.

It has been suggested previously that the UJT may have evolved in the early multi-cellular organism as a primitive adaptation in response to a specific evolutionary pressure, namely, the need accommodate epithelial migration, plasticity, or development under the physiological constraint of preserving epithelial continuity, integrity and barrier function^25^. Observations reported here suggest that the jammed phase might be seen as a further adaptation of the confluent epithelial collective that favors metabolic economy while operating under those same physiological constraints. The unjammed phase would then be seen as an adaptation that accommodates the confluent epithelial collective to episodic dynamic events –including pattern formation, growth, remodeling, plasticity, and wound healing– but comes at elevated metabolic expense. Adaptation of energy metabolism to physiological constraints, as hypothesized here, is reminiscent of the behavior of smooth muscle, wherein the latch state is seen as the adaptation of the contractile machinery of the cell to the evolutionary pressure of minimizing metabolic energy expense under the constraint of maintaining physiological levels of isometric tone in hollow organs^65–68^ (Supplement 2). So too, this hypothesis concerning epithelial jamming and unjamming proposes that the epithelial collective has become well adapted to –and specialized for– metabolic economy.

## Acknowledgements

This work was funded by the National Cancer Institute (NCI grant number U01CA202123), the National Heart Lung and Blood Institute (NHLBI grant numbers P01HL120839 and T32 HL007118), the Department of Defense (DoD grant W81XWH-15-1-0070), the Parker B. Francis Foundation, the Lemann Foundation, Santander Universidades, Fundação Faculdade de Medicina, and the Harvard-FMUSP (Universidade de Sao Paulo) program. This work was conducted with support from Harvard Catalyst | The Harvard Clinical and Translational Science Center (National Center for Advancing Translational Sciences, National Institutes of Health Award UL 1TR002541) and financial contributions from Harvard University and its affiliated academic healthcare centers. The content is solely the responsibility of the authors and does not necessarily represent the official views of Harvard Catalyst, Harvard University and its affiliated academic healthcare centers, or the National Institutes of Health. We acknowledge the use of the facilities of the Boston University Neurophotonics Center. We gratefully acknowledge Ichun A. Chen (Boston University) and Karissa Tilbury (University of Maine) for expert assistance with FLIM, and Dr. Jonathan Dreyfuss and Harvard Catalyst for expert consultation on statistical analysis.

## Author Contributions

S.J.D., V.M.K.T., J.F. designed the study; S.J.D., V. M.K.T., J.F., J.T.G., N.C.O. performed experiments; S.J.D., J.F., S.K., C.Y.P, M.D. analyzed data; S.J.D., V. M.K.T., J.F., S.T.W., J.M., J.J.F. interpreted data; S.J.D., J.F., J.J.F. wrote the manuscript; S.J.D., V. M.K.T., J.F., J.T.G., D.R., M.H.Z., J.M., J.P.B., J.J.F. reviewed and edited the manuscript. J.J.F oversaw the project.

## METHODS

### MDCKII Peredox Cell Line

A stable cell line was made from MDCKII wild-type cells (ECACC 85011435 purchased from Sigma MTOX1300-1VL) using Lipofectamine 3000 transfection of the Peredox^30,69^ construct (pcDNA3.1-Peredox-mCherry-NLS, AddGene 32384). As it is widely considered that no diffusion barriers exist between the cytoplasmic and nuclear compartments^70^, we regard the redox ratio measured the Peredox NLS construct to be indicative of the redox ratio in the cytoplasm. Additionally, the NLS tag allows us to distinguish individual cells during multi-channel fluorescence microscopy by facilitating cell segmentation. Transfection is followed by a period of G418 (Geneticin) selection, FACS sorting, and monoclonal expansion. MDCKII cells were expanded, passed, and cultured in DMEM (Gibco) supplemented with 5.6 mM glucose, 4mM glutamine, 110 mg/L sodium pyruvate and phenol red, 5% Fetal bovine serum (FBS) (Atlanta Biologicals), 100 U/mL Penicillin, and 100 μg/mL Streptomycin (Gibco). Cells were cultured in incubators maintained at 37°C and at 5% CO_2_. Cells are cultured in T_75_ flasks in 13 mL of culture media. Medium was exchanged once 24 hrs after each passage and then every second day thereafter. Cells were passed once they reached 70-80% confluency.

### Polyacrylamide Gels

We use polyacrylamide (PA) gels with a stiffness of 9.6 kPa^71^. Gels are cast in 12-well glass bottom plates. Each well is treated with 1 mL of bind silane solution (80 μL acetic acid glacial, 50 μL 3-(trimethoxysilyl)propyl methacrylate, and 200 mL deionized water) for 1 hour at room temperature. Glass bottom plates were then rinsed thoroughly with distilled water and dried with an air gun. The polyacrylamide gel consisted of 188 μL of a 40% acrylamide solution, 59 μL of a 2% bisacrylamide solution, 0.5 μL TEMED, 100 μL of 5% ammonium persulfate (APS) solution, and 650 μL ultrapure water. A volume of 24 μL of the polyacrylamide mixture was pipetted into each well of the glass bottom plate and subsequently covered with an 18 mm glass coverslip to uniformly spread the liquid across the bottom of the well resulting in a 100 µm thick gel. After the gel was completely polymerized, DI water was added to each well to prevent gels from drying out. The glass coverslips were then removed.

### Collagen-1 and Micro-Bead Coating

Polymerized polyacrylamide gels were first treated with 440 μL of a 1:50 (v/v) mixture of Sulfosuccinimidyl 6-(4'-azido-2'-nitrophenylamino)hexanoate (Sulfo-SANPAH) (ProteoChem) and 50 mM HEPES solution under a UV light for 10 minutes. Afterwards, gels were rinsed with HEPES buffer followed by 3 rinses with PBS. 150 μL of fluorescent carboxyl polystyrene microspheres (Glacial Blue, 0.2 μm diameter, Bangs Labs FCGB003) diluted at 1:6000 in DI water was applied to each gel for 30 minutes and then subsequently washed 5x with PBS. Bovine Collagen-1 (Advanced BioMatrix) stock solution was diluted in HEPES buffer solution to make a 0.2 mg/mL collagen solution. This solution was filtered with a 0.2 μm Acrodisc syringe filter (Pall). Immediately after filtering, 1 mL of collagen diluted solution was pipetted onto each gel. The plate was covered and placed in a 4°C fridge overnight. The following day, collagen was aspirated and rinsed twice with HEPES buffer solution.

### PDMS barrier preparation

Large, thin Polydimethylsiloxane (PDMS) layers were made by mixing 8 g Silicone Elastomer Base with 0.8 mg Silicone Elastomer curing agent (Sylgard® 184 silicone elastomer kit). Mixture was spread in a 150×15 petri dish (VWR) and degassed for 30 minutes. Petri dishes were placed in an oven to cure overnight at 65°C. PDMS blanks were cut from the resulting film using an 18 mm diameter hollow punch. In the center of each circular PDMS disk, a 3×12 mm rectangle was cut by hand using a scalpel. Cut PDMS masks were put in 70% ethanol overnight. Following the ethanol treatment, PDMS masks were treated with a 1% BSA (Sigma) and 1% Pluronic (Sigma) mixture overnight in a plate shaker set to low speed, and subsequently rinsed 5 times with PBS and air dried before use^72^. Immediately prior to cell seeding, gels and PDMS stencils were sterilized by exposure to UV light for 10 minutes before placing them in the biosafety hood. PDMS stencils were briefly air dried in the biosafety hood. The PBS in the wells was aspirated and PDMS stencils were placed on the gels.

### Cell redox experiments

80,000 cells suspended in 50 μL of culture media were seeded on the exposed collagen-1 coated polyacrylamide gel surface within each 3×12 mm PDMS mask region. After 2 hours, wells were filled with 3 mL of culture media. Culture Media was exchanged every subsequent day before measurement. Cells were grown for two days after seeding until dense, confluent layers were formed before lifting PDMS barriers. PDMS barriers were removed with tweezers at timepoints 24 hours and 4 hours prior to imaging. The staggard removal of the barriers in time allowed for all conditions to be imaged simultaneously and ensured that all cells were in culture for identical amounts of time before data acquisition. Two hours prior to the Peredox bioprobe measurements, cell culture medium was replaced with media containing no sodium pyruvate in order to reduce effects of exogenous pyruvate or lactate excreted by the cells on the redox state. This imaging media contained no phenol red to minimize background during fluorescence microscopy imaging. Following this media exchange, cells were moved onto a fluorescent microscope stage housed within an incubator set to 37°C and 5% CO_2_ to stabilize fluctuations in redox prior to imaging. Cells were imaged on a Leica (DMi8) using a 20x objective at 2048×2048 pixels and with a resolution of 0.325 µm/pixel. We used a Texas Red fluorescence cube to image the Peredox mCherry fluorophore and a custom filter (Chroma filter set: ET405/20x, T425lpxr, ET525/50m) to image the Peredox t-Sapphire fluorophore. Cell boundaries are also imaged with phase contrast microscopy and glacial blue beads were imaged with a DAPI channel. Images in all four channels were acquired across the narrow (3 mm) axis of the cell layer and at the center of the long (12 mm) axis of the cell layer from edge-to-edge with 10% imaging overlap, thus capturing roughly 20,000 cells per layer. After each experiment, the Peredox bioprobe was calibrated as outlined in Hung *et al*^69^. Medium containing varying amounts of lactate and pyruvate (Supplementary Table 2) were added to each well, incubated for 8 minutes, and imaged in the Texas Red and t-Sapphire channels, resulting in approximately 15 minutes between successive calibration images. The calibration media contains no glucose and no phenol red. After the final calibration step, wells were rinsed with PBS. Then, 350 μL of Trypsin (Corning) with 10% Triton X-100 (Sigma) was pipetted onto the surface of each gel and allowed to sit for 5 minutes in order to delaminate cells from the collagen-1 coated PA gel surface. Final bead positions were acquired using the same fluorescence imaging channels as the initial data measurement. Redox experiments were repeated on three separate occasions with each repeat containing four biological replicates per condition. Here, biological replicates are considered individual wells that contain an epithelial cell layer prepared in a manner as described above. Each replicate was only measured once as described above. We subsequently pooled data of single-cell measurements from all repeats and replicates to generate graphical data. The analysis of the pre-barrier lift condition contained a total of n = 53,826 cells, the 4-hour condition contained a total of n = 129,455 cells, and the 24-hour condition contained a total of n = 156,866 cells.

### Glucose uptake experiment

MDCKII wild-type cells were seeded and cultured in collagen-1 coated polyacrylamide gels in 12-well glass bottom plates and PDMS barriers were removed as described above. Glucose uptake was measured by the addition of 400 µM 2-deoxy-2-[(7-nitro-2,1,3-benzoxadiazol-4-yl)amino]-D-glucose (2-NBDG, Cayman Chemical Company 11046) diluted in 2 mL of glucose-free DMEM to each well. After one hour of incubation, cells were rinsed with 1 ml of warm DPBS and fixed with 4% PFA for 20 minutes at room temperature. Cells were rinsed 3x with PBS and refrigerated for 24 hours. The following day, cells were imaged on a Leica (DMi8) fluorescent microscope using a GFP channel and 20x objective. Glucose uptake experiments were repeated twice, with two biological replicates for each condition. In one preparation, each biological replicate was imaged at two different regions per cell layer, and the other replicate was imaged at four different regions per cell layer for a combined total of six measurements per condition.

### Mitochondrial membrane potential experiments

MDCKII wild-type cells were seeded and cultured in collagen-1 coated polyacrylamide gels in 12-well glass bottom plates and PDMS barriers were removed as described above for the Peredox redox experiment. Cells were treated with a spike of 25 nM Tetramethylrhodamine ethyl ester perchlorate (TMRE, Sigma 87917) for 30 minutes before imaging. A negative control was performed on a subset of the wells by treating cells with a spike of 10 µM Carbonyl cyanide 4-(trifluoromethoxy)phenylhydrazone (FCCP, Abcam ab120081) 10 minutes prior to the application of TMRE. Cells were imaged on a Leica (DMi8) fluorescent microscope using a Texas Red channel and 20x objective. Immediately following TMRE imaging, cells were rinsed with 1 ml of warm DPBS and fixed with 4% PFA for 20 minutes at room temperature. Cells were then rinsed 3x with PBS. After fixation, 200 nM MitoView-Green (Biotium 70054) diluted in PBS was added to each well and left to incubate for 2 hours. MitoView-Green is a membrane potential independent dye and is used here to assess whether mitochondrial mass varies across the cell layer. After incubation, MitoView-Green is imaged as before using a GFP filter cube. Mitochondrial membrane potential experiments were completed once with three biological replicates. Each replicate was imaged at four different regions per cell layer.

### Fluorescence Lifetime Imaging (FLIM)

MDCKII wild-type cells were seeded and cultured in collagen-1 coated polyacrylamide gels inside 35 mm glass bottom petri dishes and PDMS barriers were removed as described above. FLIM on expanding monolayers was carried out using a multiphoton microscope (Ultima Investigator, Bruker) equipped with a plate heater (TC-324B, Warner Instruments) and an objective heater (TC-HLS-05, Bioscience Tools) to maintain the sample at 37°C for the duration of the imaging study. A titanium:sapphire laser (InSight X3, Spectra-Physics) was tuned to 76o nm for two-photon excitation of NADH. The laser beam was focused onto the sample through a 16x water-immersion objective (Nikon, 0.8 N.A., 3 mm working distance). The fluorescence emission from NADH was transmitted through a dichroic mirror (700 nm, short-pass) that separates the excitation and emission paths, passed through a clean-up filter (720 nm, short-pass), and was collected through a bandpass filter (440/40 nm, band-pass) to a photon counting photomultiplier tube (10770PB-50, Hamamatsu). Images were collected using 1024 × 1024 pixels at a resolution of 0.792 µm/pixel and a pixel dwell time of 4 µs. Tiled z-stacks were acquired to capture a 811 µm wide strip of cells from edge to edge, with the number of tiles (4 to 7, using a 2% overlap) and slices (4 to 9, in 5 µm steps) that were determined depending on the width and on the inclination of each monolayer. Time-correlated single photon counting electronics (SPC-150, Becker&Hickl) was set to 60 seconds collection time and 256 time bins were used to acquire the temporal decay of NADH fluorescence. Second-harmonic generated signals from self-assembled collagen fibers^73^ (760 nm excitation, 375/30 nm emission) were used to measure the instrument response function (IRF), which had a full width at half maximum of 311 ps. FLIM experiments contain three biological replicates. Each replicate was imaged at two different regions per cell layer. As such, each imaging region provides data corresponding to two migrating edges (one moving leftward, and one moving rightward). Graphical data is therefore generated by pooling all 12 images for each condition.

### Peredox Cell Analysis

All images are first flat field corrected using the ImageJ BaSiC plugin^74^. Using the ImageJ BaSiC plugin, fluorescence images are additionally dark-field corrected. Images were registered using custom MATLAB (R2017b Mathworks, Natick, MA) software to account for microscope stage positioning error when computing cell nuclei displacements. All analysis steps were completed on individual imaging fields-of-view and resulting data was reconstructed into a contiguous ‘stitched’ dataset after image analysis was completed. Quantities measured on a per cell basis (cell speed, tractions, shape, area, and cytoplasmic redox potential) are linearly interpolated onto a grid with grid spacing chosen to be consistent with typical cell spacing in the dense layer (35 pixels or ~11 µm). Composite colormaps are then generated by computing the average x-position of cells (i.e., the center of mass of the layer) across the entire imaging region, and aligning all layers by their center x-position. After alignment, the gridded quantities are then averaged in a pixel-by-pixel fashion. Traces are generated by binning data along the entire y-direction as well as in bins along the x-direction of approximately 260 µm in width. Error bars represent layer-to-layer standard deviation of the means.

For dynamical measurements, we use the sequence of images collected during the Peredox cell calibration in which images are recorded once every 15 minutes for a total of 8 frames. During this image sequence, nuclei positions, trajectories, and segmentations are measured from the mCherry fluorescence channel of the Peredox sensor using custom MATLAB analysis software which we call tracking with dynamic segmentation (TWDS). This algorithm uses optical flow to estimate the motion of nuclei from one frame to the next, which is combined with segmentation of the nuclei to determine the new position of the nucleus. Accordingly, if in subsequent frames, overlapping nuclei are not properly segmented, the two overlapping nuclei can be distinguished by TWDS based upon their previous position history and their interpolated position as given by optical flow. Finally, care was taken in the implementation of TWDS not to rely on intensity thresholding to identify nuclei. From the tracked nuclei data, cell speeds are computed from inter-frame displacements representing the magnitude of the velocity vector only. Cell morphology is derived using built-in MATLAB region-property functions of a Voronoi tessellation generated from the nuclei centroid positions.

Cell tractions are measured using images of the Glacial Blue labeled microbeads. Bead image pairs consist of one image taken during the Peredox redox measurement and one image taken after the Peredox calibration and cell trypsinization are completed. Image pairs are registered with custom MATLAB code to correct for microscope stage positioning error which occurs as a result of repeated removal of the multi-well plate for media exchanges. Bead displacements are measured using optical flow. From bead displacement measurements, cell tractions are calculated as previously described^28^. Using the cell segmentation information outlined above, we extract the traction forces beneath each individual cell footprint and calculate the average magnitude of the tractions exerted by each cell.

Redox ratio: The Peredox biosensor was analyzed on a per-cell basis (Supplementary Fig. 1). Using the nuclei tracking and segmentation outlined above, the average intensity of pixels associated with each cell nucleus was calculated separately for both the mCherry (red) and T-Sapphire (green) fluorescence channels for every image over the 8 image sequence (1 data measurement plus 7 calibration steps outlined above). The ratio of the red-to-green fluorescence signal (proportional to NAD+/NADH) was then computed for each cell in each image in the sequence. To compute a fluorescence response and calibration curve, use the cells imaged in a single field-of-view at the leading edge of a 24 hr timepoint, and we follow steps and fitting procedures as outlined by Hung *et al*^69^. Briefly, we fit the fluorescence response obtained during the lactate-pyruvate titration to a logistic function:

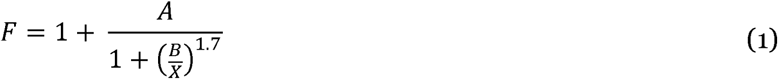

Where *F* is the normalized fluorescence ratio of the Peredox red-to-green channels for each cell, *X* is the lactate-to-pyruvate ratio used in the titration, and A and B are fit parameters. We use a Hill Coefficient of 1.7 as characterized elsewhere^69^. After fitting the normalized fluorescence response of each cell within a single image at the leading edge of the advancing cell layer at the 24 hr timepoint to the logistic function using MATLAB, we find average fit parameters of *A* = 1.04 (±0.24) and *B* = 44.13 (±26.08). Next to generate the calibration curve, which we use to convert the fluorescence measurement of the Peredox bioprobe to the NAD+/NADH ratio, we apply the method which uses the LDH equilibrium^31^.

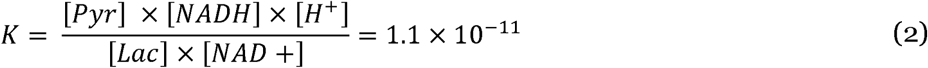

In equation (2), *K* is the equilibrium constant for LDH. We assume a physiological pH of 7.4 and thus [H^+^] = 10^−7.4^. Using the logistic fit parameters from the leading edge of the 24 hr timepoint, we solve equations (1) and (2) for the NAD+/NADH ratio resulting in a fluorescence-ratio-to-redox-ratio calibration.

All Peredox datasets were combined while in the fluorescence (red/green) ratio format into average composite images or x-profile traces of the fluorescence ratio. After the composites were generated, we converted the fluorescence ratio to NAD+/NADH using the calibration as outlined above. To compute error bars, we measured the standard deviation of the mean fluorescence ratio (red/green) of each sample. The standard deviation was then added or subtracted to the mean ratio, and then this resulting fluorescence ratio was transformed to NAD+/NADH ratio using the calibration method outlined above.

### Analysis of 2-NBDG and TMRE Images

Prior to analysis of glucose uptake (2-NBDG) and mitochondrial membrane potential (TMRE) images, all images are flat field corrected using the ImageJ BaSiC plugin^74^. Fluorescence images are additionally dark-field corrected to reduce background. Images were stitched together by concatenating adjacent fields-of-view into long strips which span the epithelial cell layer from edge-to-edge. For the 2-NBDG and TMRE images, binary masks which distinguish the free-space where no cells exist from epithelial cells within the layer were generated by hand using ImageJ. The center of each layer was then found by computing the center-of-mass of the cell layer based on the masked area. Each imaged strip across the cell layer thus results in two advancing fronts which were treated as two separate layers. Composite colormaps are then generated by aligning all layers by their center x-position with the advancing layer moving in the positive X-direction. After alignment, images are then averaged in a pixel-by-pixel fashion. Traces are generated by binning data along the entire y-direction as well as in bins along the x-direction of approximately 260 µm in width. Error bars represent layer-to-layer standard deviation of the means.

The 2-NBDG data was normalized by cell area in order to capture the amount of 2-NBDG, and hence glucose uptake, in each cell as opposed to each pixel as shown in the fluorescence image. The cell area normalization curve comes from the average cell areas measured from the Peredox cell experiments which is the same data as shown in the composite colormaps of figure 2.

MitoView analysis followed the same procedure as the 2-NBDG and TMRE data with the exception of the masking step. Images from the MitoView tended to have extensive large and saturating bright puncta throughout the layer indicative of aggregated dye or non-specific binding to particulate debris left over from the PDMS barrier. To mask these regions and omit them from analysis, we used Ilastik, the interactive learning and segmentation toolkit^75^. After this masking step, analysis followed as before.

### FLIM Image Analysis

Binary masks were generated by analyzing stitched NADH intensity images using Ilastik, the interactive learning and segmentation toolkit^75^. To isolate signals from cytoplasmic NADH, we set four pixel classification categories (i.e., cell cytoplasm, cell nuclei, debris/remnant PDMS, and background). Using these pixel classifiers, we generated individual masks for all z-slices. We then projected all pixels from slices that contained nuclei and debris into the other slices. This process resulted in a 3D binary mask for each FLIM stack that identifies the cell cytoplasm while omitting cell nuclei, debris, and background (Supplementary Fig. 2). These binary masks were imported in MATLAB (R2019a, Mathworks, Natick, MA) and used to isolate and analyze lifetime data only from the cell cytoplasm. For each pixel in the stitched image stack, time decays of fluorescence intensity from different z-slices were added to create a 2D FLIM data set that incorporates 3D spatial information. Binning lifetime data in both space^73^ (i.e., adding photon counts from a moving window of 2o pixels on each side of the current pixel) and time^76^ (i.e., adding photon counts from 4 consecutive time bins to achieve a total of 64 time bins) was implemented to maximize the total photon count in the temporal decays used for further analysis^77^. Lifetime images resulting from combining unbinned Ilastik masks and binned lifetime data yield lower variability and higher goodness-of-fit in the lifetime estimates (Supplementary Fig. 2). Temporal decays of fluorescence intensity were described via a two-exponential decay model:

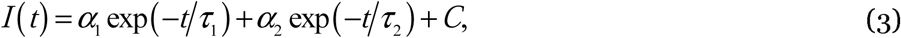

where *C* represents a constant level of background, while *τ*_1_ and *τ*_2_ represent, respectively, the fast and slow rates of decay with fractional contributions *α*_1_ and *α*_2_ (with *α*_1_ + *α*_2_ = 1). It should be noted that the fast and slow rates of decay represent, respectively, the lifetimes of free and enzyme-bound NADH^36^. For each pixel, the assumed decay (Equation 3) was convolved with the experimentally measured IRF and compared with the binned temporal decay curve. Nonlinear least square analysis was performed using the MATLAB built-in function lsqnonlin and assigning random initial guesses and physical bounds to the lifetime parameters. Values of the fractional contributions and lifetimes were constrained so that 0 ≤ *α*_*i*_ ≤ 1, while 0.02 ns ≤ *τ*_*i*_ ≤ 100 ns^73^. Upon convergence, the goodness-of-fit was calculated using the coefficient of determination R^2^ ∈ [0,1] and the mean fluorescence lifetime (*τ*_*m*_) was computed as a weighted average of the fast and slow lifetimes: *τ*_*m*_ = *α*_1_*τ*_1_ + *α*_2_*τ*_2_. The regression algorithm was parallelized to handle large FLIM data sets. A two-exponential decay model fits NADH lifetime data well and reveals spatial heterogeneities in the binding state of NADH across the migrating epithelial monolayer (Supplementary Fig. 3). The above analysis was validated by measuring the fluorescent lifetimes of Rhodamine B (Sigma 83689) in distilled water (10^−2^M). Our custom analysis led to lifetime values (*τ*_1_ = 0.11±0.003, *τ*_2_ = 0.70±0.01) consistent with those obtained using the commercial software SPCImage (*τ*_1_ = 0.16±0.09, *τ*_2_ = 0.70±0.10) and with previously published data^78^.

## Data Availability

The datasets generated during and/or analyzed during the current study are available from the corresponding author on reasonable request.

## REFERENCES

1. Bernstein, B. W. & Bamburg, J. R. Actin-ATP Hydrolysis Is a Major Energy Drain for Neurons. Journal of Neuroscience 23, 1–6 (2003).

2. Daniel, J. L., Molish, I. R., Robkin, L. & Holmsen, H. Nucleotide exchange between cytosolic ATP and F-actin-bound ADP may be a major energy-utilizing process in unstimulated platelets. European Journal of Biochemistry 156, 677–683 (1986).

3. Bhattacharya, D., Azambuja, A. P. & Simoes-Costa, M. Metabolic Reprogramming Promotes Neural Crest Migration via Yap/Tead Signaling. Developmental Cell 53, 199-211.e6 (2020).

4. Shiraishi, T. et al. Glycolysis is the primary bioenergetic pathway for cell motility and cytoskeletal remodeling in human prostate and breast cancer cells. Oncotarget 6, 130–143 (2014).

5. Beckner, M. E., Stracke, M. L., Liotta, L. A. & Schiffmann, E. Glycolysis as Primary Energy Source in Tumor Cell Chemotaxis. J Natl Cancer Inst 82, 1836–1840 (1990).

6. Gatenby, R. A. & Gillies, R. J. Why do cancers have high aerobic glycolysis? Nature Reviews Cancer 4, 891–899 (2004).

7. Humphries, B. A. et al. Plasminogen Activator Inhibitor 1 (PAI1) Promotes Actin Cytoskeleton Reorganization and Glycolytic Metabolism in Triple-Negative Breast Cancer. Mol Cancer Res 17, 1142–1154 (2019).

8. Zhang, J. et al. Energetic regulation of coordinated leader–follower dynamics during collective invasion of breast cancer cells. PNAS 116, 7867–7872 (2019).

9. Hanahan, D. & Weinberg, R. A. Hallmarks of cancer: the next generation. Cell 144, 646–74 (2011).

10. Warburg, O., Wind, F. & Negelein, E. The Metabolism of Tumors in the Body. The Journal of General Physiology 8, 519–530 (1927).

11. Epstein, T., Gatenby, R. A. & Brown, J. S. The Warburg effect as an adaptation of cancer cells to rapid fluctuations in energy demand. PLOS ONE 12, e0185085 (2017).

12. Lu, J. The Warburg metabolism fuels tumor metastasis. Cancer Metastasis Rev 38, 157–164 (2019).

13. Im, M. J. C. & Hoopes, J. E. Energy metabolism in healing skin wounds. Journal of Surgical Research 10, 459–464 (1970).

14. Jones, J. D., Ramser, H. E., Woessner, A. E., Veves, A. & Quinn, K. P. Quantifying Age-Related Changes in Skin Wound Metabolism Using In Vivo Multiphoton Microscopy. Advances in Wound Care 9, 90–102 (2019).

15. Li, J. et al. Investigating the healing mechanisms of an angiogenesis-promoting topical treatment for diabetic wounds using multimodal microscopy. Journal of Biophotonics 11, e201700195 (2018).

16. Bays, J. L., Campbell, H. K., Heidema, C., Sebbagh, M. & DeMali, K. A. Linking E-cadherin mechanotransduction to cell metabolism through force-mediated activation of AMPK. Nat Cell Biol 19, 724–731 (2017).

17. Bartolák-Suki, E. & Suki, B. Tuning mitochondrial structure and function to criticality by fluctuation-driven mechanotransduction. Scientific Reports 10, 1–13 (2020).

18. Hu, H. et al. Phosphoinositide 3-Kinase Regulates Glycolysis through Mobilization of Aldolase from the Actin Cytoskeleton. Cell 164, 433–446 (2016).

19. Atia, L. et al. Geometric constraints during epithelial jamming. Nature Physics 14, 613–620 (2018).

20. Mongera, A. et al. A fluid-to-solid jamming transition underlies vertebrate body axis elongation. Nature 561, 401–405 (2018).

21. Angelini, T. E. et al. Glass-like dynamics of collective cell migration. Proc Natl Acad Sci U S A 108, 4714–9 (2011).

22. Haeger, A., Krause, M., Wolf, K. & Friedl, P. Cell jamming: collective invasion of mesenchymal tumor cells imposed by tissue confinement. Biochim Biophys Acta 1840, 2386–95 (2014).

23. Oswald, L., Grosser, S., Smith, D. M. & Käs, J. A. Jamming transitions in cancer. J. Phys. D: Appl. Phys. 50, 483001 (2017).

24. Park, J. A. et al. Unjamming and cell shape in the asthmatic airway epithelium. Nat Mater 14, 1040–8 (2015).

25. Mitchel, J. A. et al. The unjamming transition is distinct from the epithelial-to-mesenchymal transition. Nature Communications (in press).

26. Kılıç, A. et al. Mechanical forces induce an asthma gene signature in healthy airway epithelial cells. Scientific Reports 10, 966 (2020).

27. Bi, D., Lopez, J. H., Schwarz, J. M. & Manning, M. L. A density-independent rigidity transition in biological tissues. Nature Physics 11, 1074–1079 (2015).

28. Trepat, X. et al. Physical forces during collective cell migration. Nature Physics 5, 426–430 (2009).

29. Serra-Picamal, X. et al. Mechanical waves during tissue expansion. Nature Physics 8, 628–634 (2012).

30. Hung, Y. P., Albeck, J. G., Tantama, M. & Yellen, G. Imaging cytosolic NADH-NAD(+) redox state with a genetically encoded fluorescent biosensor. Cell Metab 14, 545–54 (2011).

31. Williamson, D. H., Lund, P. & Krebs, H. A. The redox state of free nicotinamide-adenine dinucleotide in the cytoplasm and mitochondria of rat liver. Biochem J 103, 514–27 (1967).

32. Reffay, M. et al. Interplay of RhoA and mechanical forces in collective cell migration driven by leader cells. Nature Cell Biology 16, 217–223 (2014).

33. Ladoux, B. & Mège, R.-M. Mechanobiology of collective cell behaviours. Nature Reviews Molecular Cell Biology 18, 743–757 (2017).

34. Hino, N. et al. ERK-Mediated Mechanochemical Waves Direct Collective Cell Polarization. Developmental Cell 53, 646–660.e8 (2020).

35. Chance, B., Schoener, B., Oshino, R., Itshak, F. & Nakase, Y. Oxidation-reduction ratio studies of mitochondria in freeze-trapped samples. NADH and flavoprotein fluorescence signals. J. Biol. Chem. 254, 4764–4771 (1979).

36. Lakowicz, J. R., Szmacinski, H., Nowaczyk, K. & Johnson, M. L. Fluorescence lifetime imaging of free and protein-bound NADH. PNAS 89, 1271–1275 (1992).

37. Becker, W. Fluorescence lifetime imaging – techniques and applications. Journal of Microscopy 247, 119–136 (2012).

38. Becker, W. Introduction to Multi-dimensional TCSPC. in Advanced Time-Correlated Single Photon Counting Applications (ed. Becker, W.) 1–63 (Springer International Publishing, 2015). doi:10.1007/978-3-319-14929-5_1.

39. Bird, D. K. et al. Metabolic Mapping of MCF10 A Human Breast Cells via Multiphoton Fluorescence Lifetime Imaging of the Coenzyme NADH. Cancer Res 65, 8766–8773 (2005).

40. Drozdowicz-Tomsia, K. et al. Multiphoton fluorescence lifetime imaging microscopy reveals free-to-bound NADH ratio changes associated with metabolic inhibition. JBO 19, 086016 (2014).

41. Yaseen, M. A. et al. Fluorescence lifetime microscopy of NADH distinguishes alterations in cerebral metabolism in vivo. Biomed. Opt. Express, BOE 8, 2368–2385 (2017).

42. Sharick, J. T. et al. Protein-bound NAD(P)H Lifetime is Sensitive to Multiple Fates of Glucose Carbon. Scientific Reports 8, 5456 (2018).

43. Stringari, C. et al. Metabolic trajectory of cellular differentiation in small intestine by Phasor Fluorescence Lifetime Microscopy of NADH. Scientific Reports 2, 568 (2012).

44. Stringari, C., Nourse, J. L., Flanagan, L. A. & Gratton, E. Phasor Fluorescence Lifetime Microscopy of Free and Protein-Bound NADH Reveals Neural Stem Cell Differentiation Potential. PLOS ONE 7, e48014 (2012).

45. Zou, C., Wang, Y. & Shen, Z. 2-NBDG as a fluorescent indicator for direct glucose uptake measurement. J Biochem Biophys Methods 64, 207–15 (2005).

46. O’Neil, R. G., Wu, L. & Mullani, N. Uptake of a Fluorescent Deoxyglucose Analog (2-NBDG) in Tumor Cells. Mol Imaging Biol 7, 388–392 (2005).

47. Yamada, K., Saito, M., Matsuoka, H. & Inagaki, N. A real-time method of imaging glucose uptake in single, living mammalian cells. Nat Protoc 2, 753–62 (2007).

48. Perry, S. W., Norman, J. P., Barbieri, J., Brown, E. B. & Gelbard, H. A. Mitochondrial membrane potential probes and the proton gradient: a practical usage guide. Biotechniques 50, 98–115 (2011).

49. Spurlin, J. W. et al. Mesenchymal proteases and tissue fluidity remodel the extracellular matrix during airway epithelial branching in the embryonic avian lung. Development 146, dev175257 (2019).

50. Kim, J. H. et al. Unjamming and collective migration in MCF10A breast cancer cell lines. Biochemical and Biophysical Research Communications 521, 706–715 (2020).

51. Palamidessi, A. et al. Unjamming overcomes kinetic and proliferation arrest in terminally differentiated cells and promotes collective motility of carcinoma. Nature Materials 18, 1252–1263 (2019).

52. Staneva, R. et al. Cancer cells in the tumor core exhibit spatially coordinated migration patterns. J Cell Sci 132, (2019).

53. Veerati, P. C. et al. Airway mechanical compression: its role in asthma pathogenesis and progression. European Respiratory Review 29, (2020).

54. Streichan, S. J., Hoerner, C. R., Schneidt, T., Holzer, D. & Hufnagel, L. Spatial constraints control cell proliferation in tissues. PNAS 111, 5586–5591 (2014).

55. Vander Heiden, M. G., Cantley, L. C. & Thompson, C. B. Understanding the Warburg effect: the metabolic requirements of cell proliferation. Science 324, 1029–33 (2009).

56. Zanotelli, M. R. et al. Regulation of ATP utilization during metastatic cell migration by collagen architecture. MBoC 29, 1–9 (2018).

57. Le, A. et al. Inhibition of lactate dehydrogenase A induces oxidative stress and inhibits tumor progression. PNAS 107, 2037–2042 (2010).

58. Angelini, T. E., Hannezo, E., Trepat, X., Fredberg, J. J. & Weitz, D. A. Cell migration driven by cooperative substrate deformation patterns. Phys Rev Lett 104, 168104 (2010).

59. Sun, F., Dai, C., Xie, J. & Hu, X. Biochemical issues in estimation of cytosolic free NAD/NADH ratio. PLoS One 7, e34525 (2012).

60. Blacker, T. S. & Duchen, M. R. Investigating mitochondrial redox state using NADH and NADPH autofluorescence. Free Radic Biol Med 100, 53–65 (2016).

61. DeMarzio, M. et al. Genomic signatures of the unjamming transition in compressed human bronchial epithelial cells. BioRxiv.

62. Salvi, A. M. & DeMali, K. A. Mechanisms linking mechanotransduction and cell metabolism. Curr Opin Cell Biol 54, 114–120 (2018).

63. Tambe, D. T. et al. Collective cell guidance by cooperative intercellular forces. Nat Mater 10, 469–75 (2011).

64. Krishnan, R. et al. Fluidization, resolidification, and reorientation of the endothelial cell in response to slow tidal stretches. Am J Physiol Cell Physiol 303, C368–C375 (2012).

65. Dillon, P. F., Aksoy, M. O., Driska, S. P. & Murphy, R. A. Myosin phosphorylation and the cross-bridge cycle in arterial smooth muscle. Science 211, 495–497 (1981).

66. Hai, C. M. & Murphy, R. A. Regulation of shortening velocity by cross-bridge phosphorylation in smooth muscle. American Journal of Physiology-Cell Physiology 255, C86–C94 (1988).

67. Murphy, R. A. What is special about smooth muscle? The significance of covalent crossbridge regulation. The FASEB Journal 8, 311–318 (1994).

68. Fredberg, J. J. et al. Friction in airway smooth muscle: mechanism, latch, and implications in asthma. J Appl Physiol (1985) 81, 2703–12 (1996).

69. Hung, Y. P. & Yellen, G. Live-cell imaging of cytosolic NADH-NAD+ redox state using a genetically encoded fluorescent biosensor. Methods Mol Biol 1071, 83–95 (2014).

70. Zhang, Q., Piston, D. W. & Goodman, R. H. Regulation of Corepressor Function by Nuclear NADH. Science 295, 1895–1897 (2002).

71. Mih, J. D. et al. A multiwell platform for studying stiffness-dependent cell biology. PLoS One 6, e19929 (2011).

72. Sunyer, R. et al. Collective cell durotaxis emerges from long-range intercellular force transmission. Science 353, 1157–61 (2016).

73. Becker, W. The bh TCSPC Handbook 8th ed. Becker & Hickl GmbH https://www.becker-hickl.com/literature/handbooks/the-bh-tcspc-handbook/(2019).

74. Peng, T. et al. A BaSiC tool for background and shading correction of optical microscopy images. Nature Communications 8, 14836 (2017).

75. Berg, S. et al. ilastik: interactive machine learning for (bio)image analysis. Nature Methods 16, 1226–1232 (2019).

76. Walsh, A. J., Sharick, J. T., Skala, M. C. & Beier, H. T. Temporal binning of time-correlated single photon counting data improves exponential decay fits and imaging speed. Biomed Opt Express 7, 1385–1399 (2016).

77. Datta, R., Heaster, T. M., Sharick, J. T., Gillette, A. A. & Skala, M. C. Fluorescence lifetime imaging microscopy: fundamentals and advances in instrumentation, analysis, and applications. J Biomed Opt 25, (2020).

78. Kristoffersen, A. S., Erga, S. R., Hamre, B. & Frette, Ø. Testing Fluorescence Lifetime Standards using Two-Photon Excitation and Time-Domain Instrumentation: Rhodamine B, Coumarin 6 and Lucifer Yellow. J Fluoresc 24, 1015–1024 (2014).

