## Supplement for "Epithelial layer unjamming shifts energy metabolism toward glycolysis"

1. Department of Environmental Health, Harvard T.H. Chan School of Public Health, Boston, MA, USA; 2. Faculdade de Medicina FMUSP, Universidade de Sao Paulo, Sao Paulo, SP, BR; 3. Department of Biomedical Engineering, Boston University, Boston, MA, USA; 4. Howard Hughes Medical Institute, Boston University, Boston, MA, USA; 5. Department of Medicine, Brigham and Women’s Hospital and Harvard Medical School, Boston, MA, USA.

**Supplement 1:**

**Cell migration, metabolism, and the ECM.** In the recent literature, references to cell metabolism and mechanics typically focusses upon ECM mechanics rather than cell/tissue mechanics. It is well established that mechanical properties of the extracellular matrix (ECM) in the tumor microenvironment dramatically affect cancer cell migration and invasion and hence the events that lead to metastasis^1–7^. With regards to metabolism, recent reports have demonstrated that cell energy metabolism is highly mechanoresponsive to cell-ECM interactions, yet with few predictable trends^6–11^. For example, increasing matrix stiffness has been shown to increase ATP/ADP ratios and increase mitochondrial respiration while also increasing the invasiveness of pancreatic ductal adenocarcinoma^7^. In contrast, in transformed non-small-cell human bronchial epithelial cells, higher matrix stiffness has been shown to increase glycolytic rates through actomyosin-matrix coupling^10^. To further illustrate this unpredictable response to ECM, in highly invasive triple-negative breast cancer cells, decreasing matrix stiffness has been shown to increase oxidative phosphorylation while in contrast, MCF10A cells increase their rate of glycolysis. Meanwhile, less invasive MCF-7 and T-47D cells show little metabolic change regardless of substrate^8^. It is clear that cancer cell metabolism and migration is highly dependent on the microenvironment, however, the story of cancer cell migration, invasion, and metabolism, through the lens of ECM is not clear.

**Supplement 2:**

**Metabolic economy.** The concept of metabolic economy was first introduced into the literature in the context of muscle biology^12–15^. Richard Murphy argued persuasively that striated muscle versus smooth muscle are the products of distinctly different evolutionary pressures. On the one hand, striated muscle is disposed in the body so as to perform mechanical work. Specifically, contraction of the activated striated muscle acts to create a defined mechanical displacement of some defined mechanical load, as in the acts of running across the savanna, the lifting of a weight or the pumping of blood. The ability to perform such mechanical work at the smallest metabolic expenditure offers an obvious survival advantage.

The ratio of the work performed to the metabolic energy expended for any given muscle task is called the muscle efficiency, which for maximally activated striated muscle falls close to 25%^14^. On the other hand, smooth muscle is disposed in hollow organs such as blood vessels, airways, bladder, gut, and iris. In these hollow organs smooth muscles can and do perform some mechanical work, which in many cases is central to function, as in peristalsis in the gut. But their main function is most often to maintain organ shape, size, and tone, as in airways and blood vessels. Importantly, even when mechanical work is nil, as in the case of isometric contraction, these critical functions require metabolic energy nonetheless. In such cases the muscle efficiency is not a particularly useful metric.

A more useful metric, for example, might be the rate of metabolic energy expenditure that is required to maintain a fixed level of isometric tensile stress. Murphy called this metric the muscle economy, as distinct from and not be confused with muscle efficiency. In its economy smooth muscle is seen to excel. For example, to attain some given level of isometric tensile stress, smooth muscle is more economical than striated muscle by a factor of 300. Generation of the same tensile stress at only a miniscule fraction of the energy expenditure is an astounding muscle adaptation. This remarkable economical state, known as the ‘latch state’, is attributable in large part to the overall down-regulation of acto-myosin cycling rates in smooth compared with striated muscle, together with the unique capability during isometric contraction for smooth muscle to further down-regulate its acto-myosin cycling rates and plastically remodel its cytoskeleton^16,17^. When smooth muscle does shorten or lengthen to a different operating length, as in micturition, it upregulates its actomyosin cycling rate. It switches from slowly cycling latch bridges to rapidly cycling cross bridges, but does so at the cost of an increased rate of energy utilization. When the new operating length is attained the cycling rates are downregulated, latch is re-established, and energy utilization is diminished. Perhaps an even more remarkable example of economy is the catch state of the mollusk hinge muscle, wherein the rates of actomyosin cycling and metabolic energy utilization can go to zero. Striated muscle is well adapted for energetic efficiency whereas smooth muscle is well adapted for energetic economy.

**Supplement 3: Local cell perimeter is the strongest predictor of local cell migration speed and local NAD+/NADH ratio.**

Tendencies noted in Figs. 1 and 2 in the main text, as well systematic relationships between morphological, mechanical and metabolic indices, are analyzed here statistically and quantified on a cell-by-cell basis. We have pooled data from 138,973 cells from 12 wells measured at the 24 hr timepoint, 122,277 cells from 7 wells at the 4hr timepoint and 51,001 cells from 3 wells at no Lift condition. To find the strongest statistical predictors of local cell migration speed as well as local NAD+/NADH ratio, we used LASSO (least absolute shrinkage and selection operator) regression^18^, which is available as the open-source glmnet package^19^ for Matlab (<https://glmnet.stanford.edu/>). First, to normalize skewed distributions we converted all mechanical, morphological and metabolic indices (aspect ratio, area, perimeter, shape index, traction magnitude, speed and Peredox fluorescent ratio) to a log scale^20^ and then standardized all indices to have mean of 0 and standard deviation of 1. We then ran LASSO regression for migration speed as the dependent variable with all the other mechanical, morphological, and metabolic indices as input variables. With a lambda value that minimized the mean cross-validated error, we established a linear regression equation for migration speed, $Y= \sum_{i=1}^{n} {(\beta}_{i}X_{i})+c$ with the coefficients for the input variables listed in the table below. Entries of zero indicate variables that, based on the data, LASSO did not select as being meaningful, and thus constrained to zero.

At the 24hr timepoint, we found that while controlling for the other variables in the model, local cell perimeter had the largest coefficient and thus was the strongest predictor of local migration speed. X-position was the second strongest predictor and the Peredox fluorescent ratio (NAD+/NADH) was the 7^th^. Interestingly, the coefficients for cell area and nuclear perimeter were not selected by LASSO regression and therefore constrained to be 0. This could be because these indices are not independent of cell perimeter and LASSO regression is known to choose one among several covariates^19^. At the 4hr timepoint, we found that the mean squared error increased but nevertheless, local cell perimeter was still the strongest predictor of local migration speed. At the No-Lift condition, X-position and local cell traction were the 2 strongest predictors.

Next, we ran LASSO regression with the Peredox fluorescent ratio (NAD+/NADH) as the dependent variable with all the other indices (including migration speed as input variables). Overall, the regression was less accurate than the ones for migration speed. Nevertheless, we again found that the absolute coefficient of local cell perimeter was the biggest and thus the strongest statistical predictor at both 24hr and 4hr timepoints.

Supplementary Table 1: Coefficients for the indices for LASSO regression

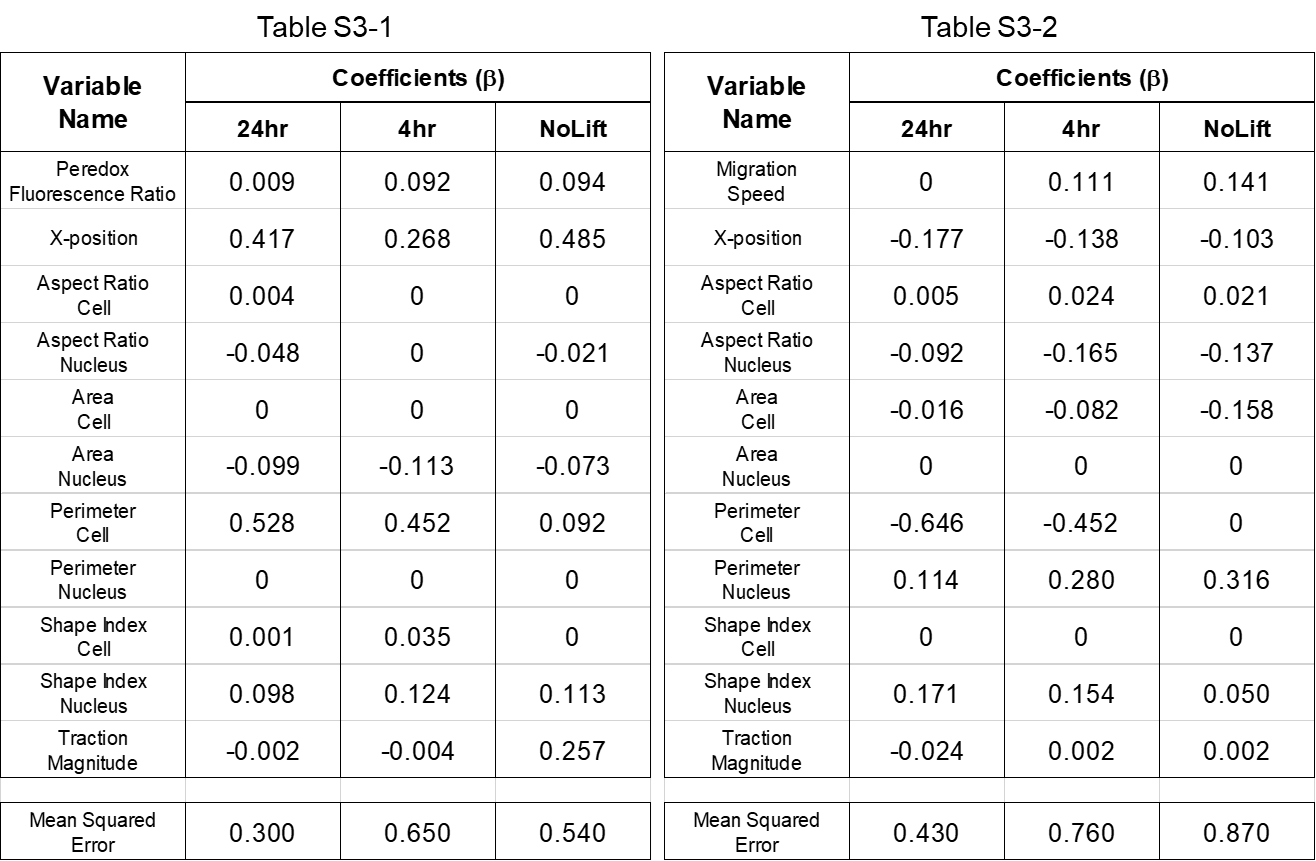

**Supplementary Movie 1: Cells confined by the PDMS barrier remain non-motile, solid-like, and jammed.**

This movie shows MDCKII Peredox cells confined by the PDMS barrier in phase imaging microscopy and fluorescence imaging microscopy of the same region. Cells throughout the layer show little migration and essentially no change in cell shape. The left-most side of the imaged region represents the center (bulk) of the expanding strip of cells. The right-side of the imaged region where no cells are visible is the PDMS barrier. Images were captured using a Leica (DMi8) equipped with a 20x objective. Fluorescence movies represent the red (mCherry) channel of the nuclear localized Peredox biosensor acquired using a Texas Red filter cube. All images were acquired at 2048x2048 pixels and with a resolution of 0.325 µm/pixel. Successive fields of view were imaged with 10% overlap and subsequently concatenated together using Matlab. After stitching, the image data is resized in ImageJ to reduce overall file size of the avi movie. The Peredox calibration procedure and the cell trypsinization protocol for traction microscopy are destructive endpoints to the experiment. As such, the cells shown here are separate preparations from those presented in the main text. However, the cell preparation is identical to that outlined in the methods section.

**Supplementary Movie 2: After lifting the PDMS barrier, cells unjam, increase in migration speed, elongate in shape, and migrate into the free space.**

This movie shows MDCKII Peredox cells immediately after removal of the PDMS barrier in phase imaging microscopy and fluorescence imaging microscopy of the same region. Cells near the leading edge of the advancing layer increase in migration speed and elongate in shape. As cells migrate into the free space, a wave of unjamming propagates retrograde from the leading edge into the densely packed bulk enlisting cells to begin migrating. The left-most side of the imaged region represents the center (bulk) of the expanding strip of cells. The right-side of the imaged region where no cells are visible is the free-space created after lifting the PDMS barrier. Images were captured using a Leica (DMi8) equipped with a 20x objective. Fluorescence movies represent the red (mCherry) channel of the nuclear localized Peredox biosensor acquired using a Texas Red filter cube. All images were acquired at 2048x2048 pixels and with a resolution of 0.325 µm/pixel. Successive fields of view were imaged with 10% overlap and subsequently concatenated together using Matlab. After stitching, the image data is resized in ImageJ to reduce overall file size of the avi movie. The Peredox calibration procedure and the cell trypsinization protocol for traction microscopy are destructive endpoints to the experiment. As such, the cells shown here are separate preparations from those presented in the main text. However, the cell preparation is identical to that outlined in the methods section.

**Supplementary Table 2: Peredox cell calibration media lactate and pyruvate concentrations.**

| **Calibration Medium** | **Lactate (mM)** | **Pyruvate(mM)** |
| --- | --- | --- |
| **A** | 20 | 0 |
| **B** | 20 | 0.04 |
| **C** | 20 | 0.14 |
| **D** | 20 | 0.4 |
| **E** | 20 | 1 |
| **F** | 20 | 3.34 |
| **G** | 0 | 20 |

**Supplementary Figure 1**

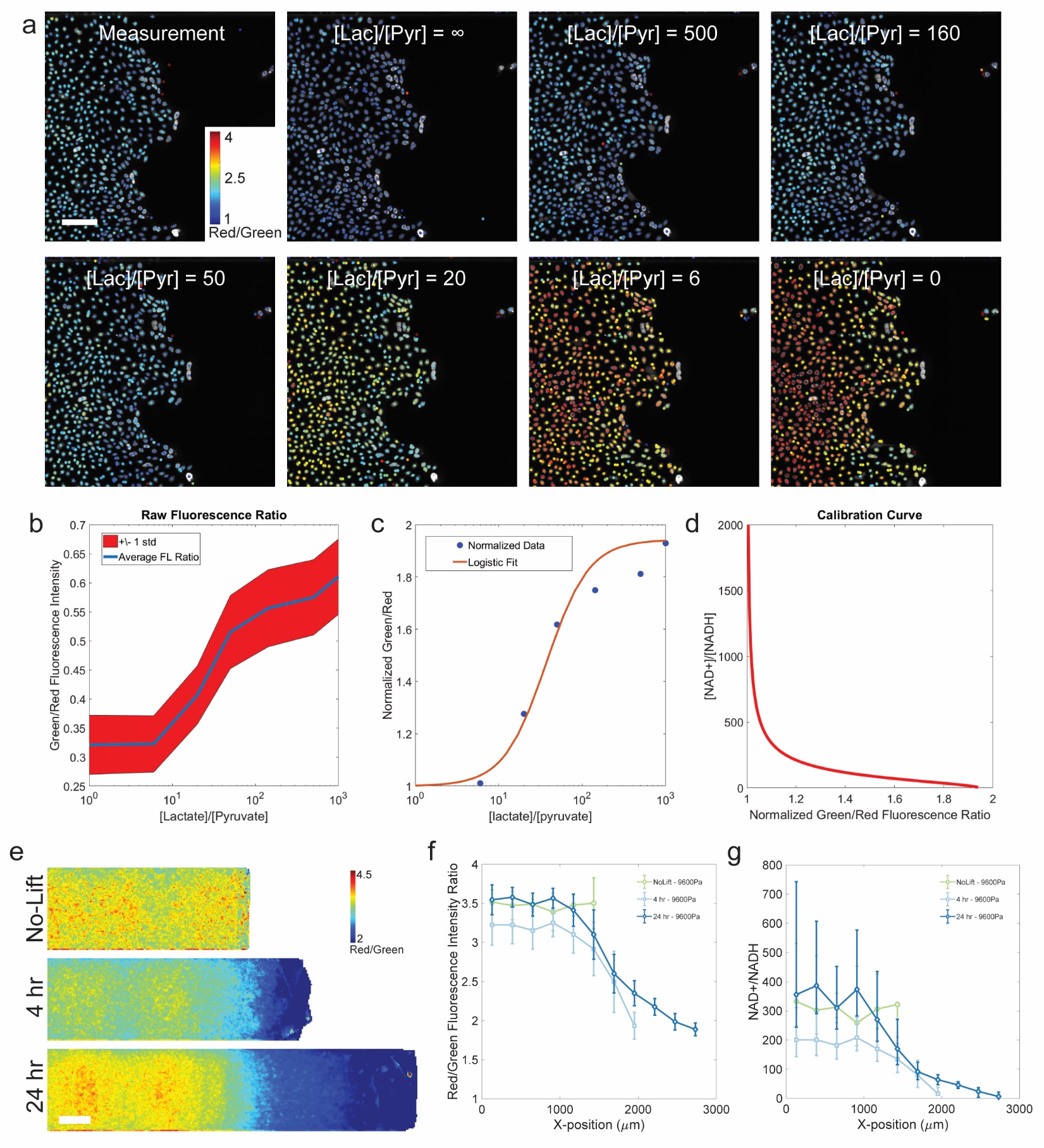

**LEGEND: Peredox redox biosensor calibration.**

**a** We use a field of view from the leading edge of an advancing MDCKII cell layer at the 24 hr timepoint to calibrate the fluorescence ratio of all samples. Shown here is a sequence of images of the red nuclei channel of the Peredox biosensor starting first with the unperturbed measurement of the biosensor, followed by a series of successive images with a decreasing ratio of exogenously supplemented [lactate]/[pyruvate] (Supplementary Table 1). The color dot at the center of each cell nucleus represents the red-to-green ratio for each cell identified in our custom Matlab cell tracking and segmentation software. As the [lactate]/[pyruvate] ratio decreases, the red-to-green ratio increases as expected. **b** The average raw fluorescence ratio (green/red) of the Peredox biosensor of all cells in the image for each of the [lactate]/[pyruvate] conditions. The red shaded region shows the standard deviation of the fluorescence ratio above and below the curve at each titration point. **c** The average fluorescence response curve is normalized as outlined in Hung *et al* and fit with equation 1 shown in the Methods section^21^. The average fit parameters found by fitting each cell individually with the logistic function are A = 1.04 (±0.24) and B = 44.13 (±26.08) and are subsequently used to calculate the final calibration curve. **d** To calculate the fluorescence calibration curve, we solve the equilibrium equation of LDH (equation 2) for the NAD+/NADH ratio. **e** The average spatial map of the raw fluorescence ratio (red/green) of the Peredox biosensor is shown prior to barrier lift and at 4 hrs and 24 hrs after barrier lift. **f** The average trace of the raw fluorescence ratio (red/green) for each timepoint is generated by binning data from each cell layer along the y-axis and into 260 µm wide bins along the x-axis. Error bars represent layer-to-layer standard deviations of the mean. **g** The average trace of the fluorescent ratio is then transformed using the calibration curve to result in the calibrated NAD+/NADH ratio recapitulated here and shown in the main text.

**Supplementary Figure 2**

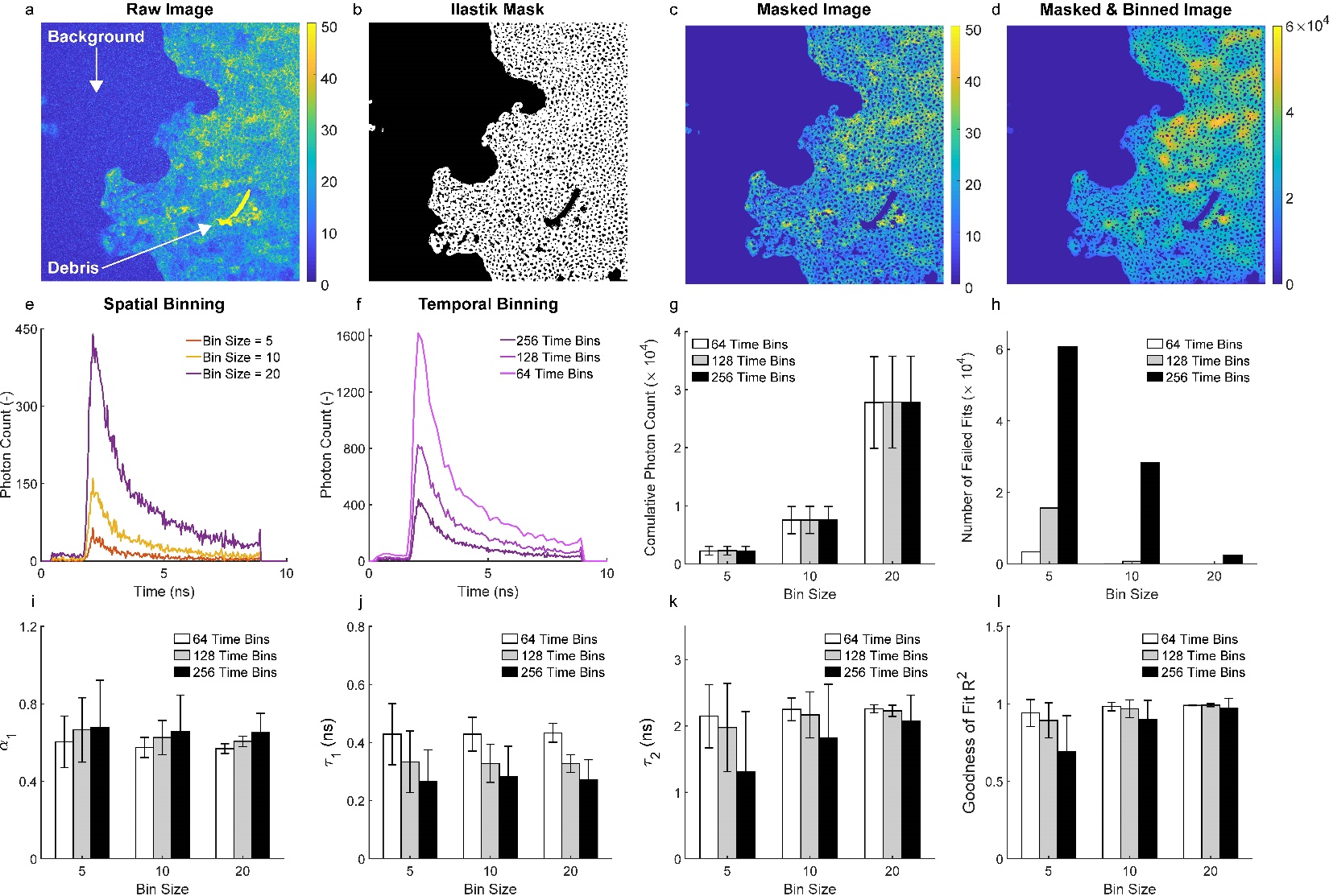

**LEGEND: FLIM image analysis masking and data binning procedure.**

**a** Raw NADH intensity image collected at one edge of the monolayer after 24 hours of expansion. NADH intensity is calculated by adding up all photons detected within a pixel during the collection time of a FLIM scan. White arrows indicate the background (low intensity signal) and debris / remnant PDMS (high intensity signal). **b** Binary mask generated using the Ilastik pixel classification module. Areas in white indicate the cell cytoplasm while areas in black indicate the cell nuclei, background, and debris that need to be removed prior to FLIM data analysis. **c** Masked NADH intensity image emphasizes cytoplasmic signals. **d** Effect of spatial binning on the NADH intensity image. By combining data from 20 adjacent pixels, one can increase dramatically the photon count in each pixel (note the difference in range between the colormaps in panels c and d). **e** Effect of spatial binning. For a representative pixel, time decays of NADH fluorescence are show for increasing spatial bin size. **f** Effect of temporal binning. For a representative pixel and a spatial bin size of 20 pixels, time decays of NADH fluorescence are shown for a decreasing number of time bins (i.e., lower temporal resolution). **g,h** Spatial binning, not temporal binning, impacts the cumulative photon count per pixel while both binning approaches lead to a reduction in the number of failed fits (lack of convergence in the curve fitting algorithm). It should be noted that, by increasing the spatial bin size and decreasing the number of time bins, we are able to fit data from all pixels within the field of view shown in panels a-to-d. **i,j,k,l** Estimated parameters of the two exponential decay model for various combinations of spatial and temporal binning. Fractional contribution of free NADH (*α_1_*), lifetimes of free and bound NADH (*τ_1_* and *τ_2_*), and goodness of fit (R^2^). Increasing the spatial bin size and decreasing the number of time bins changes the mean value of the estimated parameters while yielding a lower variability and a higher goodness of fit.

**Supplementary Figure 3**

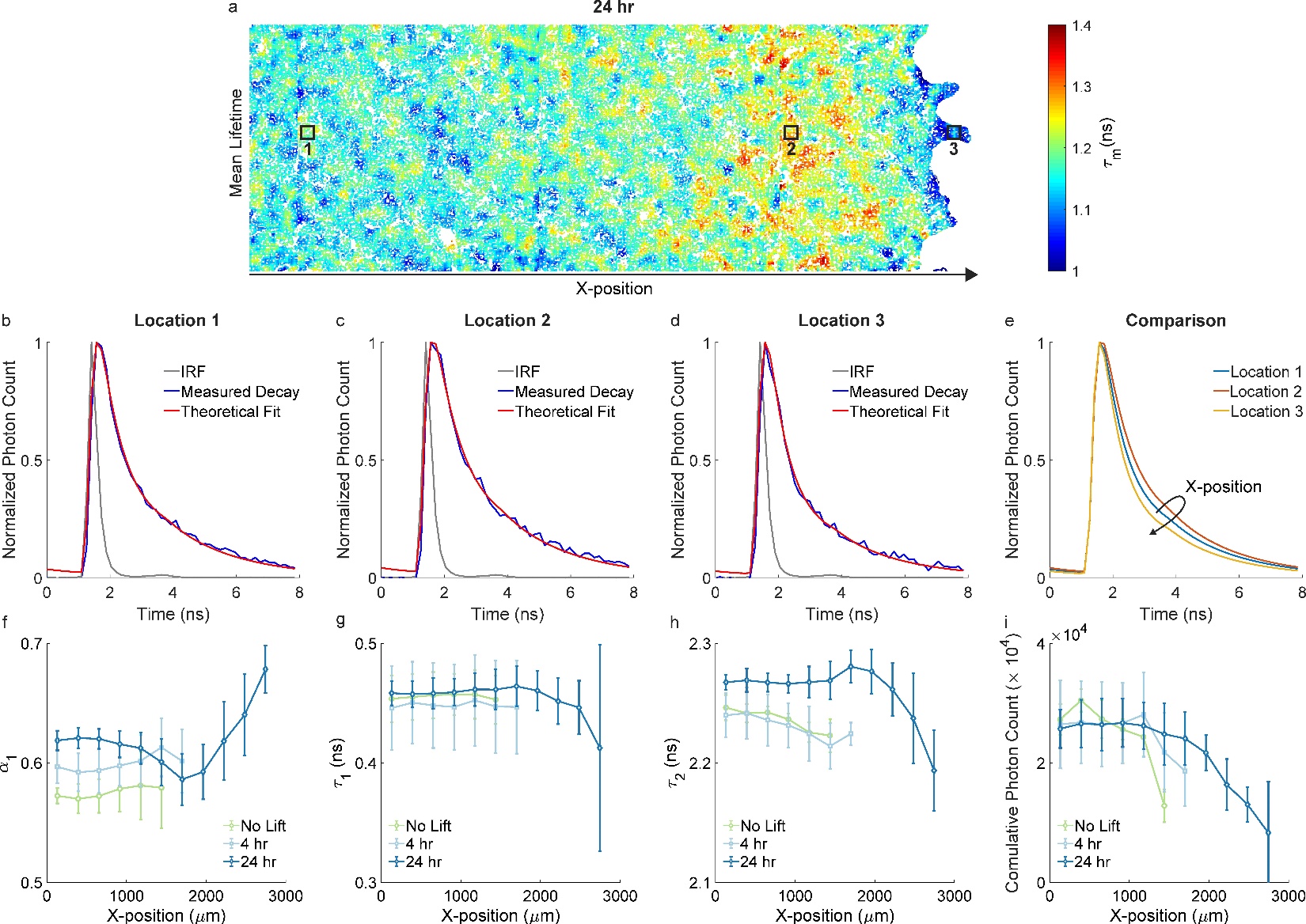

**LEGEND: Spatial variations in NADH fluorescence decays and FLIM parameters.**

**a** Spatial variation in the mean NADH lifetime (*τ_m_*) of a representative sample 24 hours after barrier removal. The FLIM colormap is obtained by using a spatial binning of 20 pixels and 64 time bins prior to curve fitting analysis. Black boxes indicate distinct locations at different X-positions along the width of the expanding monolayers. **b,c,d** Time decay of NADH fluorescence corresponding to a pixel from each of the three selected locations. The experimentally measured decay is shown in blue, the instrument response function (IRF) is shown in gray, while the theoretical fit is shown in red. **e** The complex pattern in mean NADH lifetime shown in panel a is mirrored by the fitted time decays of NADH fluorescence: starting from the bulk of the advancing cell layer (Location 1), NADH decays first undergo a right-ward shift corresponding to increasing mean lifetimes (Location 2), and finally undergo a left-ward shift corresponding to decreasing mean lifetimes at the edge of the cell layer (Location 3). **f,g,h** Traces of individual parameters of the two exponential decay model for the non-migrating cell layer (No Lift), and for the migrating cell layer at different timepoints after barrier removal (4 hr and 24 hr). While the lifetime of free NADH (*τ_1_*) shows little differences, the fractional contribution of free NADH (*α_1_*) and the lifetime of bound NADH (*τ_2_*) show differences between the various experimental groups and complex spatial patterns after 24 hours of barrier removal. The trends in *α_1_* and *τ_2_* underlie the trends in mean NADH lifetime and free to bound NADH ratio reported in Figure 3. **i** Traces of cumulative photon counts (i.e., NADH intensity) show that a decreasing trend with increasing X-position is observed in all experimental groups. Lifetime estimates (panels f-to-g) are independent from intensity and, in fact, display different trends with respect to the cumulative photon counts.

**Supplementary Figure 4**

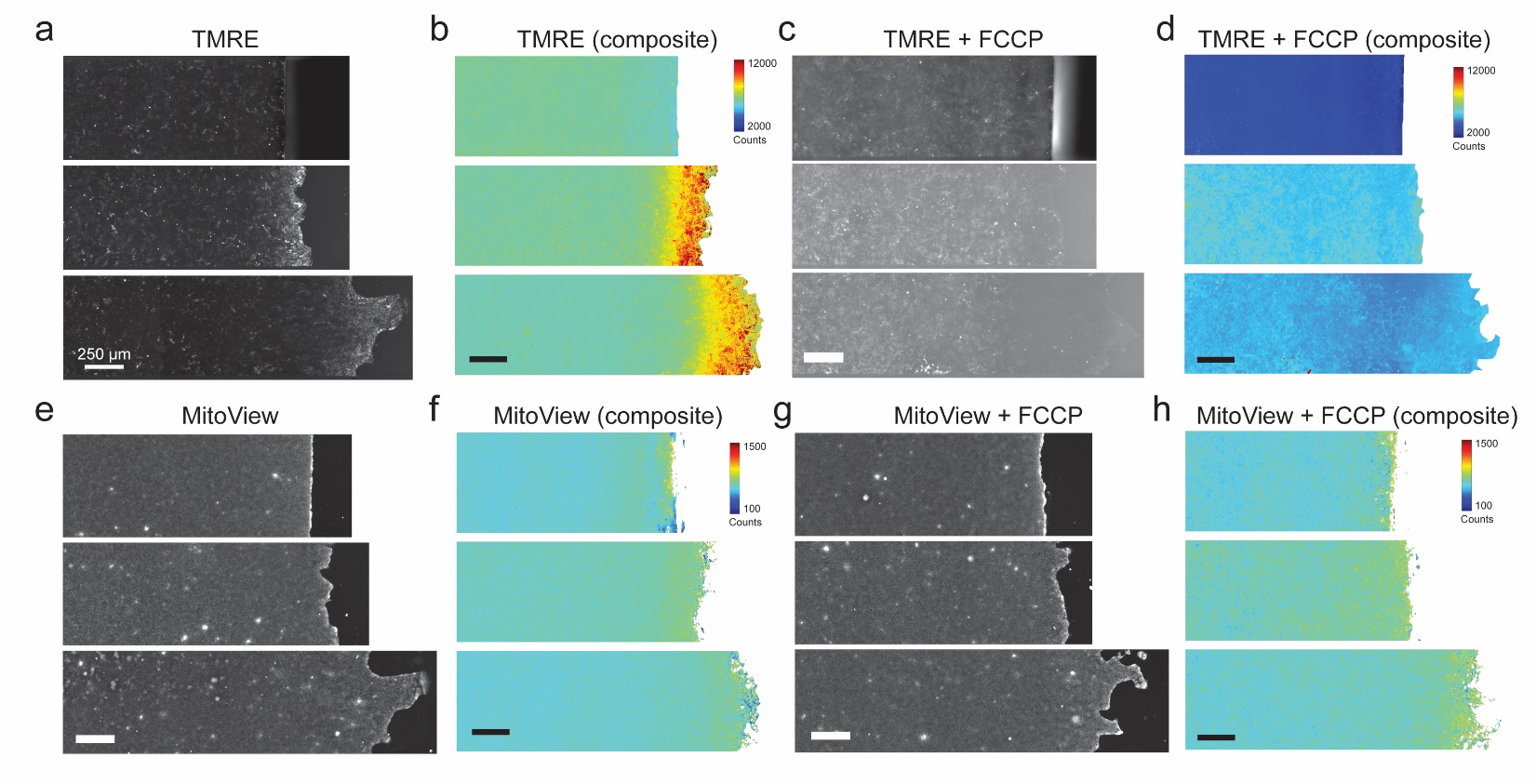

**LEGEND: Representative images and composite averages of TMRE and MitoView with and without FCCP.**

**a** Representative images of cell layers treated with TMRE, a mitochondrial membrane potential sensitive dye prior to barrier lifting and at timepoints 4 hrs and 24 hrs after barrier lifting. **b** Composite images generated from pixel-by-pixel averaging of multiple images of cell layers stained with TMRE show that prior to barrier lifting, the TMRE signal is low. At 4 hrs and 24 hrs after barrier lifting, the TMRE signal is substantially elevated only near the leading edge cells and remains low throughout the bulk. **c** Representative images of cell layers treated with TMRE and the membrane potential decoupler, FCCP prior to barrier lifting and at timepoints 4 hrs and 24 hrs after barrier lifting. **d** Composite images generated from pixel-by-pixel averaging of multiple images of cell layers stained with TMRE and treated with FCCP demonstrates substantial reductions in the TMRE signal occurs in all conditions. **e** Representative images of cell layers stained with MitoView-green, a mitochondrial membrane potential insensitive dye prior to barrier lifting and at timepoints 4 hrs and 24 hrs after barrier lifting. **f** Composite images generated from pixel-by-pixel averaging of multiple cell layers stained with MitoView show that the amount of MitoView intensity is low and uniform essentially everywhere throughout the cell layer and at all timepoints. This suggests that mitochondrial mass is uniform in all conditions and does not vary throughout the cell layer. **g** Representative images of cell layers stained with MitoView-green and treated with FCCP prior to barrier lifting and at timepoints 4 hrs and 24 hrs after barrier lifting. **g** Composite images generated from pixel-by-pixel averaging of multiple cell layers stained with MitoView and treated with FCCP show that the amount of MitoView intensity is low and uniform essentially everywhere throughout the cell layer and at all timepoints.

**Supplementary References**

1.	Alexander, N. R. *et al.* Extracellular Matrix Rigidity Promotes Invadopodia Activity. *Current Biology* **18**, 1295–1299 (2008).

2.	Artym, V. V. *et al.* Dense fibrillar collagen is a potent inducer of invadopodia via a specific signaling network. *J Cell Biol* **208**, 331–350 (2015).

3.	Paszek, M. J. *et al.* Tensional homeostasis and the malignant phenotype. *Cancer Cell* **8**, 241–254 (2005).

4.	Seewaldt, V. ECM stiffness paves the way for tumor cells. *Nature Medicine* **20**, 332–333 (2014).

5.	Wells, R. G. The role of matrix stiffness in regulating cell behavior. *Hepatology* **47**, 1394–1400 (2008).

6.	Tung, J. C. *et al.* Tumor mechanics and metabolic dysfunction. *Free Radic Biol Med* **79**, 269–80 (2015).

7.	Papalazarou, V. *et al.* The creatine–phosphagen system is mechanoresponsive in pancreatic adenocarcinoma and fuels invasion and metastasis. *Nature Metabolism* **2**, 62–80 (2020).

8.	Mah, E. J., Lefebvre, A. E. Y. T., McGahey, G. E., Yee, A. F. & Digman, M. A. Collagen density modulates triple-negative breast cancer cell metabolism through adhesion-mediated contractility. *Sci Rep* **8**, 1–11 (2018).

9.	Bartolák-Suki, E. & Suki, B. Tuning mitochondrial structure and function to criticality by fluctuation-driven mechanotransduction. *Scientific Reports* **10**, 1–13 (2020).

10.	Park, J. S. *et al.* Mechanical regulation of glycolysis via cytoskeleton architecture. *Nature* **578**, 621–626 (2020).

11.	Zanotelli, M. R. *et al.* Regulation of ATP utilization during metastatic cell migration by collagen architecture. *MBoC* **29**, 1–9 (2018).

12.	Dillon, P. F., Aksoy, M. O., Driska, S. P. & Murphy, R. A. Myosin phosphorylation and the cross-bridge cycle in arterial smooth muscle. *Science* **211**, 495–497 (1981).

13.	Hai, C. M. & Murphy, R. A. Regulation of shortening velocity by cross-bridge phosphorylation in smooth muscle. *American Journal of Physiology-Cell Physiology* **255**, C86–C94 (1988).

14.	Murphy, R. A. What is special about smooth muscle? The significance of covalent crossbridge regulation. *The FASEB Journal* **8**, 311–318 (1994).

15.	Fredberg, J. J. *et al.* Friction in airway smooth muscle: mechanism, latch, and implications in asthma. *J Appl Physiol (1985)* **81**, 2703–12 (1996).

16.	Seow, C. Y. & An, S. S. The Force Awakens in the Cytoskeleton: The Saga of a Shape-Shifter. *Am J Respir Cell Mol Biol* **62**, 550–551 (2020).

17.	Pratusevich, V. R., Seow, C. Y. & Ford, L. E. Plasticity in canine airway smooth muscle. *J Gen Physiol* **105**, 73–94 (1995).

18.	Tibshirani, R. Regression Shrinkage and Selection via the Lasso. *Journal of the Royal Statistical Society. Series B (Methodological)* **58**, 267–288 (1996).

19.	Friedman, J., Hastie, T. & Tibshirani, R. Regularization Paths for Generalized Linear Models via Coordinate Descent. *J Stat Softw* **33**, 1–22 (2010).

20.	Atia, L. *et al.* Geometric constraints during epithelial jamming. *Nature Physics* **14**, 613–620 (2018).

21.	Hung, Y. P. & Yellen, G. Live-cell imaging of cytosolic NADH-NAD+ redox state using a genetically encoded fluorescent biosensor. *Methods Mol Biol* **1071**, 83–95 (2014).
